# Deep mutational scanning of rabies glycoprotein defines mutational constraint and antibody-escape mutations

**DOI:** 10.1101/2024.12.17.628970

**Authors:** Arjun K. Aditham, Caelan E. Radford, Caleb R. Carr, Naveen Jasti, Neil P. King, Jesse D. Bloom

## Abstract

Rabies virus causes nearly 60,000 human deaths annually. Antibodies that target the rabies glycoprotein (G) are being developed as post-exposure prophylactics, but mutations in G can render such antibodies ineffective. Here, we use pseudovirus deep mutational scanning to measure how all single amino-acid mutations to G affect cell entry and neutralization by a panel of antibodies. These measurements identify sites critical for rabies G’s function, and define constrained regions that are attractive epitopes for clinical antibodies, including at the apex and base of the protein. We provide complete maps of escape mutations for eight monoclonal antibodies, including some in clinical use or development. Escape mutations for most antibodies are present in some natural rabies strains. Overall, this work provides comprehensive information on the functional and antigenic effects of G mutations that can help inform development of stabilized vaccine antigens and antibodies that are resilient to rabies genetic variation.

## Introduction

Rabies is a lyssavirus that causes encephalitis that is fatal upon symptom onset in humans and most other mammalian species^1^. The virus is common in several mammalian species^2–4^, and zoonotic human infections are responsible for ∼60,000 human deaths annually, with the greatest toll in Africa and Asia^5^. Exposure to rabies requires immediate post-exposure prophylaxis, which includes human or equine serum-derived rabies immunoglobulin (RIG) injections, monoclonal antibodies, or repeated vaccination^7^. Although prophylaxis is over 99% effective, cost and supply shortages render treatments inaccessible in some rabies-endemic regions^8–10^. Therefore, efforts are underway to develop better rabies vaccines^11–15^ and monoclonal antibody treatments^16–19^.

The main target of both vaccines and antibodies against rabies is its glycoprotein G, which mediates receptor binding^20–22^ and viral fusion with host cells^23,24^. Pre-fusion G is a trimeric complex on the viral surface^12,25^, and undergoes pH-driven conformational changes to mediate membrane fusion^26,27^. As for other class III viral fusion proteins^28–30^, rabies G is conformationally dynamic, exchanging reversibly between its pre-fusion and post-fusion conformations^24,31^. G’s conformational dynamics pose a challenge for vaccine design: the most potent neutralizing antibodies target the pre-fusion conformation, but many antibodies elicited by immunization with unmodified G protein bind to other conformations^12,25^. Therefore, efforts are underway to stabilize pre-fusion rabies G to create improved vaccine antigens^12,25^, as well as identify potently neutralizing anti-G monoclonal antibodies^19,32^.

Another challenge is that G is quite diverse, with >10% protein sequence divergence among natural rabies strains. This genetic diversity complicates monoclonal antibody development, as natural strains have been identified with resistance to antibodies, including antibodies in clinical use or development^19,26,33–35^. Attempts are being made to develop monoclonal antibodies or antibody cocktails with increased neutralization breadth and resistance to escape^19,34^.

Here we use pseudovirus deep mutational scanning^36,37^ to measure how mutations to the G ectodomain affect cell entry and neutralization by a panel of eight monoclonal antibodies. Our experiments quantify mutational constraint across G, and provide insight into how mutations affect G’s conformational dynamics and function. We also provide complete maps of escape mutations for a variety of important antibodies, enabling us to better define their epitopes and assess the extent that escape mutations for each antibody are present among natural rabies sequences. Overall, our deep mutational scan elucidates how mutations affect the function and antigenicity of G, and provides information that can help guide the design of vaccine antigens and antibody treatments.

## Results

### A pseudovirus deep mutational scan of rabies G

To measure the effects of mutations in rabies G, we used a previously described pseudovirus deep mutational scanning platform^36,37^. This platform enables creation of a library of genotype-to-phenotype linked lentiviral particles, each of which displays a unique G mutant on its surface and encodes an identifying nucleotide barcode in its genome (Fig S1A,B). These lentiviral particles (or “pseudoviruses”) encode only one viral gene (rabies G) and so can undergo only a single cycle of cellular infection, thereby enabling the study of G mutants without requiring the generation of replicative pathogenic virus. The pooled pseudovirus library can be used to infect cells under various experimental conditions, and the ability of each G mutant to enable cellular infection in each condition can be quantified by sequencing the barcodes from infected cells (Fig 1A). This approach makes it possible to measure the effects of thousands of different mutations to G in a single experiment.

**Figure 1.**
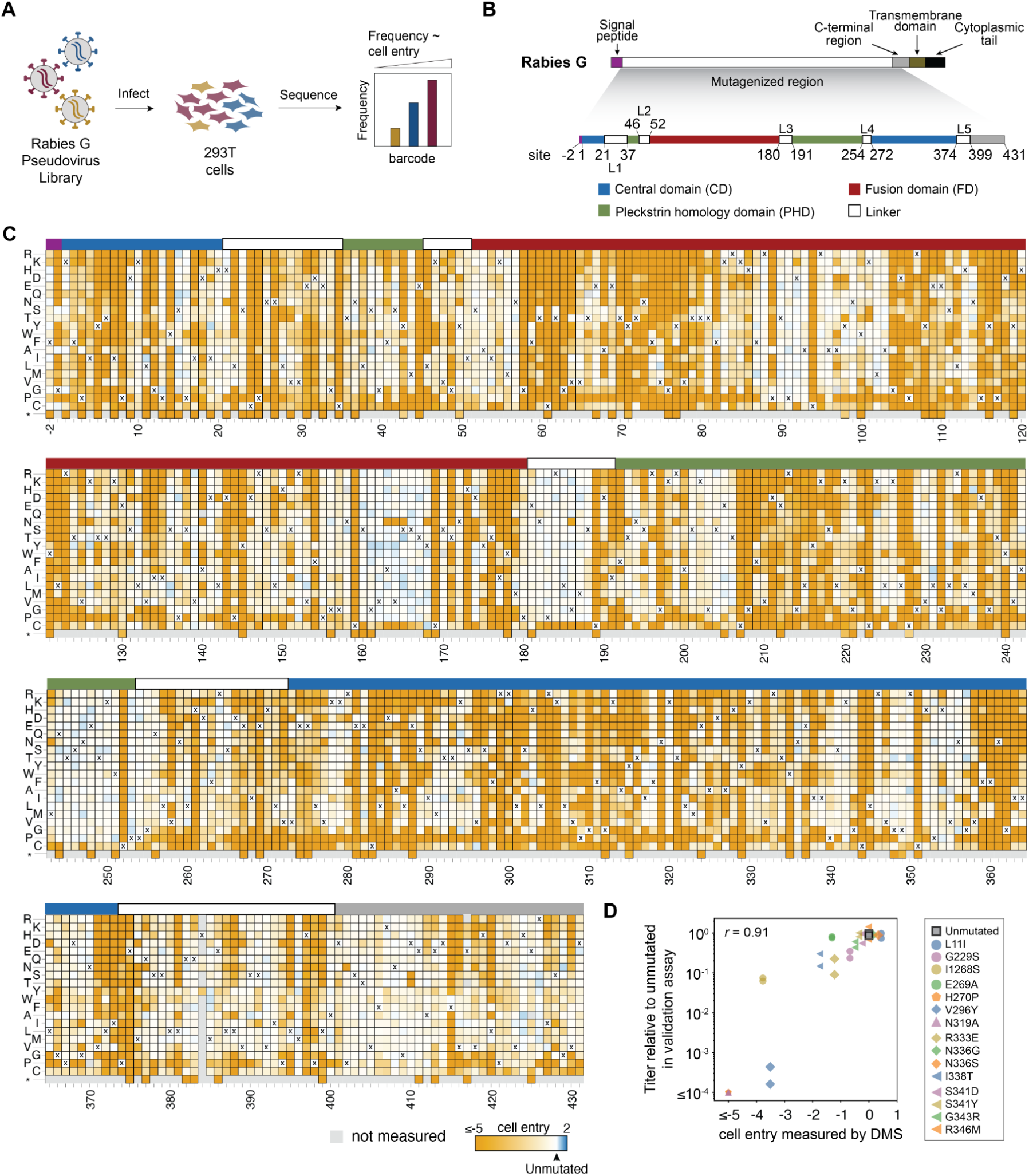
Deep mutational scanning of rabies G. (A) We create libraries of pseudoviruses expressing different mutants of G on their surface and encoding identifying nucleotide barcodes in their genomes. These libraries are used to infect cells in various conditions, and barcodes from viruses that successfully infect are deep sequenced to quantify the effects of mutations. (B) Schematic showing region of rabies G mutagenized in our libraries. We use the conventional numbering scheme where site 1 is assigned to the first site of ectodomain. In this numbering scheme, we mutagenized from site −2 to site 431. (C) Effects of mutations on G-mediated cell entry. Each column of the heatmap shows the effects of different amino acids at that site on cell entry, with the amino acid identity in the unmutated Pasteur strain G indicated with a “X”. Mutations that impair cell entry are colored orange, mutations that do not affect cell entry are white, and mutations that enhance cell entry are blue. Gray indicates mutations that were not reliably measured in the deep mutational scanning. The overlay bar indicates the regions of G. See https://dms-vep.org/RABV_Pasteur_G_DMS/cell_entry.html for an interactive version of this heatmap. (D) Validation of deep mutational scanning measurements of mutational effects on cell entry. For each of the 16 mutations (which were chosen to span a range of cell entry effects), the x-axis shows the effect measured in the deep mutational scanning (DMS) while the y-axis shows the titers of individual pseudoviruses generated with that G protein mutant relative to the unmutated parent. Two independent validation measurements of the titers were made for each mutation, hence there are two points per mutation in the plot. The *r* indicates the Pearson correlation.

We performed a deep mutational scan using the G protein from the Pasteur strain of rabies virus, a vaccine strain^15^ that recently had its pre-fusion G determined by cryo-EM^12,25^. We designed our mutant libraries to contain all single amino-acid mutations to 433 sites in G, starting two residues N-terminal to the ectodomain (at the end of the signal peptide) and continuing through the entire structurally resolved^12,25^ region of the ectodomain and into its C-terminal region (Fig 1B). The libraries therefore target 433 ⨉ 19 = 8,227 different amino-acid mutations. We also designed stop-codon mutations at 20 sites as negative controls for cell entry. Throughout this paper, we utilize the numbering scheme where one is assigned to the first site in the ectodomain, consistent with past numbering conventions for rabies G studies^12,25,26^.

We created duplicate G pseudovirus mutant libraries using previously described methods (Fig S1B)^36,37^. Each library contained ∼80,000 barcoded variants of G, and each covered >99% of the 8,227 intended ectodomain amino-acid mutations (Fig S1C). Most barcoded variants contained only one G amino-acid mutation (∼64%), although some variants contained no (∼12%) or multiple mutations (∼24%) (Fig S1D).

### Effects of mutations on G-mediated cell entry

We first measured how all G mutations affected the protein’s ability to mediate pseudovirus entry into 293T cells. We did this by infecting 293T cells with the pooled libraries of barcoded pseudovirus particles expressing the rabies G mutants, and comparing the frequency of each barcode in the infected 293T cells to a control set of cells infected with barcoded pseudovirus that display unmutated VSV-G (Fig 1A, S1B, S2A). To maximize titers of pseudovirus displaying rabies G, we performed infections with the G pseudoviruses in media supplemented with 4 µg/mL polybrene^38^ and adjusted to a pH of 7.1^39^ (Fig S2B). When pseudotyped with VSV-G, all pseudoviruses are able to enter 293T cells regardless of rabies G function, providing a baseline frequency for each barcode in the library. These baseline frequencies are then compared to the frequency of each barcode in the condition where the pseudoviruses are dependent on their rabies G mutant to infect cells. We quantify the ability of each G variant to infect cells as the logarithm of the ratio of its barcode counts relative to those for the unmutated G between the rabies-G and the VSV-G mediated infection conditions (Fig S2A). Cell entry scores greater than zero indicate improved infection relative to parental G protein, while values below zero indicate impaired entry. Since a modest fraction of barcoded variants contain multiple mutations, we employed a global epistasis model to deconvolve the effects of individual mutations^40^. Measurements of how mutations affected cell entry were highly correlated between replicates performed with the same library on different days, and between the two independent G mutant libraries (Fig S2C-D).

The effects of G mutations on cell entry are shown in Fig 1C; see the interactive displays at https://dms-vep.org/RABV_Pasteur_G_DMS/cell_entry.html to best examine the data. As expected, mutations to stop codons are uniformly deleterious for cell entry. The effects of amino-acid mutations vary widely across sites, and in some cases among different mutations at the same site. For instance, sites 160-168 in the fusion domain, 181-189 in the third linker, 201-206 and 244-250 in the pleckstrin homology domain were all tolerant of most amino-acid mutations. But other sites are highly constrained: for instance, nearly all mutations are deleterious in both fusion loops, which span sites 73-79 and 117-125^26^. In the next subsection, we discuss these patterns of constraint in more detail in the context of G’s structure.

To validate the deep mutational scanning measurements of mutation effects on cell entry, we cloned 15 individual G mutations measured to have a range of effects and determined the titers of individual pseudoviruses encoding these mutations.The effects of mutations on cell entry from the deep mutational scan were highly correlated with the individual pseudovirus titers measured in these validation assays (*r* = 0.91; Fig 1D). The deep mutational scanning is also consistent with prior experiments that have characterized the effects of individual mutations to G on its fusogenic activity^12,26^ (Fig S2E).

### Mutational constraint in the context of G’s structure and function

The effects of mutations on G’s cell-entry function varied widely across the protein, with both highly constrained and mutationally tolerant sites in all three of the central domain, pleckstrin homology domain, and fusion domain (Fig 1C, 2A). Some of this constraint can be explained relatively simply by the requirement that G fold properly in order to function. For example, one of the most constrained regions on G’s surface involves the disulfide bond connecting sites C189 and C228, where mutations to any non-cysteine amino acid at either position is highly deleterious for cell entry (Fig 2A). At sites that are buried in G’s pre-fusion structure, mutations to non-hydrophobic amino acids tend to be deleterious presumably because they disrupt the folding of protein’s hydrophobic cores (Fig S3).

**Figure 2.**
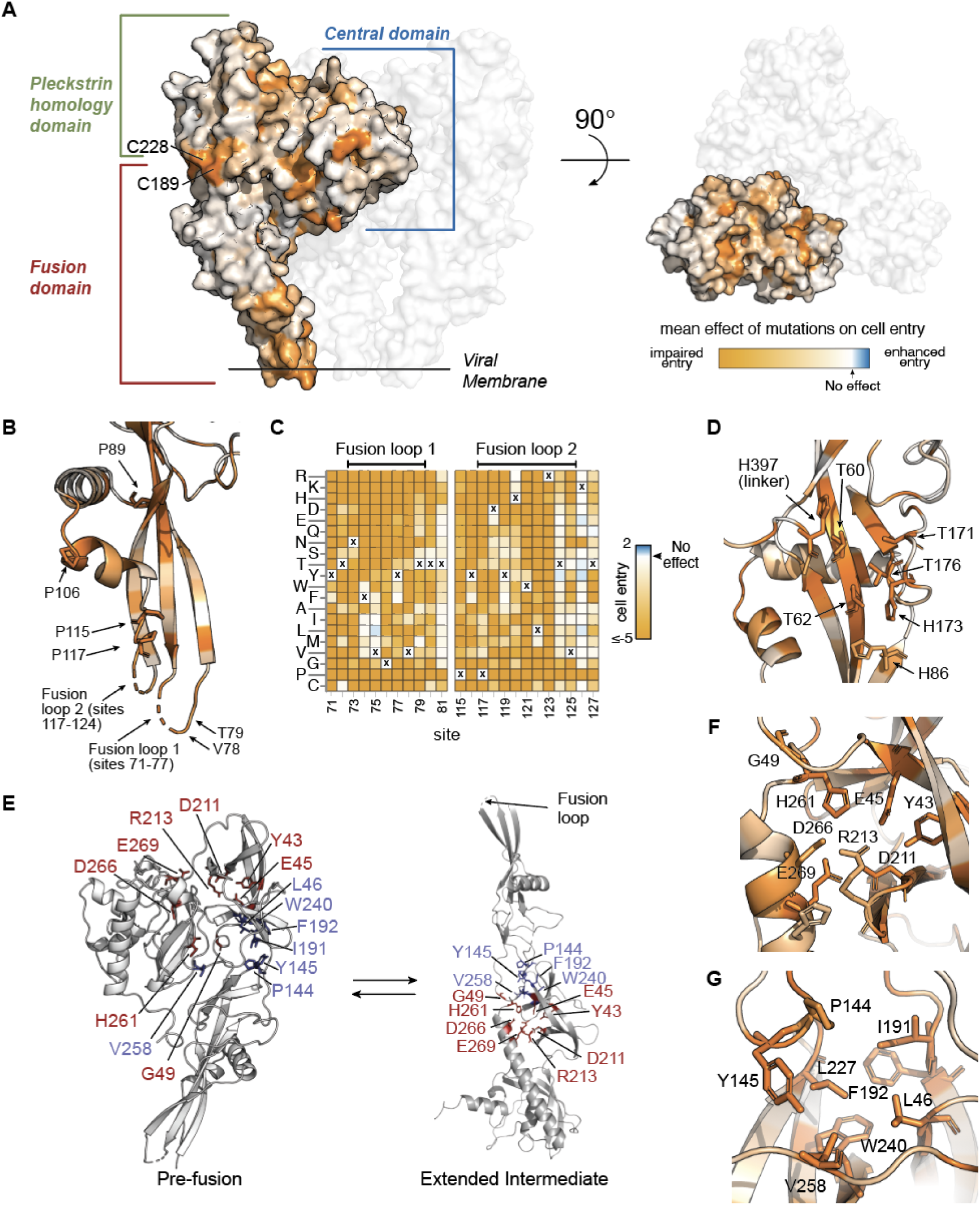
Functional constraints in the context of rabies G’s structure. (A) Pre-fusion G structure (PDB: 7U9G) with two transparent protomers and one protomer colored by mean effect of all mutations at each site on cell entry. Sites are colored orange when mutations impair cell entry, and white when they have no effect. See https://dms-vep.org/RABV_Pasteur_G_DMS/cell_entry.html#mutation-effects-on-structure-of-g for interactive structural visualizations of the effects of mutations. (B) Zoomed view of the fusion domain of a single protomer of pre-fusion G with sites colored by the mean effects of mutations on cell entry. The fusion loops are not resolved in the structure and are modeled as dashed cartoon loops. (C) Heatmaps showing the effects on cell entry of all mutations in the fusion loops and flanking sites. The parental amino-acid identities in the Pasteur strain G are indicated with “X.” (D) Zoomed view of a single protomer of pre-fusion G highlighting the histidine cluster and structurally adjacent threonines that may act as a pH sensor. Sites are colored by the mean effect mutations on cell entry as in (A). (E) A single promoter of G shown in both the pre-fusion protomer and extended intermediate conformations (PDB 6LGW), with sites highlighted in panel F shown in red and sites highlighted in panel G shown in blue. (F) Zoomed view of the extended intermediate conformation showing sites that form polar interactions in this conformation. Sites are colored by the mean effects of mutations on cell entry as in (A). (G) Zoomed view of the extended intermediate conformation showing sites that form hydrophobic interactions in this conformation. Sites are colored by the mean effects of mutations on cell entry as in (A).

Other sites of constraint provide more nuanced insights into G’s function. The fusion loops are critical to G’s cell entry mechanism, and are thought to stabilize the trimeric pre-fusion conformation of rabies G^25^. The fusion loops are only partially resolved in available three-dimensional structures of G, but our results show that many sites in both the loops and flanking beta strands are highly constrained (Fig 2B-C). Prior work has tested alanine mutations to sites F74, Y77, Y119, and W121 in the fusion loops on stability of trimeric G, and found that only Y119A does not have a strong adverse effect^25^. Consistent with this prior work, our deep mutational scan shows that a mutation to alanine is reasonably tolerated for cell entry at Y119, but not at F74, Y77 or W121 (Fig 2C). However, all of these sites except W121 do tolerate mutations to some amino acids that conserve hydrophobicity or aromaticity (e.g., F74L and Y77F are reasonably tolerated) (Fig 2C). Proline residues near the fusion loops (sites 89, 106, 115, and 117) are largely intolerant of mutations (Fig 1C, 2B), likely because these prolines are important for ensuring proper fusion loop orientation.

The deep mutational scanning also helps delineate sites that may be important in triggering the conformational change in G necessary for cell-entry. A cluster of histidines at sites 86, 173, and 397 at the interface between one of the linkers and the fusion domain is hypothesized to help govern pH-driven conformational change in rabies G^12,41^ (Fig 2D). Consistent with this hypothesis that these histidines act as a “pH sensor” crucial for G’s cell entry function, our deep mutational scan shows that mutating any of these three sites to any non-histidine amino acid impairs cell entry (Fig 1C, 2D). Our deep mutational scan also shows that threonine residues structurally adjacent to the histidines (T60, T62, T171, and T176) only tolerate serine mutations (Fig 1C, 2D), suggesting that the hydroxyl moieties of these residues may be important for G function. This finding is consistent with the possibility that these residues interact with the histidine cluster, as has been hypothesized for serines near the histidine cluster in the related VSV-G fusion protein^41^.

Our deep mutational scanning identifies some sites where the mutational constraint appears to be due to interactions in the extended intermediate conformation, which exists in equilibrium with the pre-fusion trimer^24,27,31^ (Fig 2E). A cluster of sites that includes E45, D211, R213, H261, D266, and E269 form polar and van der Waals interactions in the extended intermediate^12^ (Fig 2F), and our deep mutational scanning shows that almost all mutations at these sites are deleterious for cell entry (Fig 1C, 2F). Another cluster of sites (L46, P144, Y145, I191, F192, L227, W240, and V258) form extensive hydrophobic contacts in the extended intermediate (Fig 2E,F), and our deep mutational scanning shows that only hydrophobic mutations (or in the case of Y145, no mutations) are tolerated at these sites (Fig 1C, 2F). In the pre-fusion conformation trimer, some of these sites (eg, Y145 and V258) are distal from the other sites and largely solvent-exposed (Fig 2E), suggesting the mutational constraints originate from interactions in the extended intermediate rather than the pre-fusion conformation. These sites that stabilize the extended intermediate but likely tolerate mutation in the pre-fusion conformation represent additional candidates for stabilizing mutations that might shift the conformational equilibrium of rabies G towards the pre-fusion state.

Prior work has stabilized rabies G in its pre-fusion conformation by introducing a mutation to proline (H270P) that is tolerated in the pre-fusion conformation but is disfavored in the extended intermediate conformation^12,13^. A similar proline-based stabilization strategy has been applied to stabilize vaccine immunogens for both class I fusion proteins (eg, coronavirus spikes^42,43^) and VSV-G^44^. Consistent with the idea that a proline at site 270 in rabies G uniquely blocks conversion of the pre-fusion trimer to the extended intermediate, our deep mutational scanning shows that H270P strongly impairs cell entry but most other amino-acid mutations at this site are well tolerated (Fig S4). We used our deep mutational scanning to identify additional sites in regions that undergo large conformation changes where proline is especially disfavored (eg, K47, Q256, N259, L260, and S265) (Fig S4). These sites represent additional potential candidates for pre-fusion stabilizing proline mutations.

### Deep mutational scanning comprehensively maps antibody-escape mutations

To experimentally measure how all rabies G mutations affect antibody neutralization, we incubated the pseudovirus library with different concentrations of antibody and used deep sequencing to quantify the ability of each G variant to infect 293T cells at each antibody concentration (Fig 3A). To convert the deep-sequencing counts to neutralization values, we normalized them to the counts of a “non-neutralized standard” consisting of VSV-G pseudovirus that is not neutralized by the anti-rabies antibodies^36^ (Fig 3A). We analyzed the data using a previously described biophysical model^45^ to obtain escape values for each mutation that are proportional to the log fold-change in IC50.

**Figure 3.**
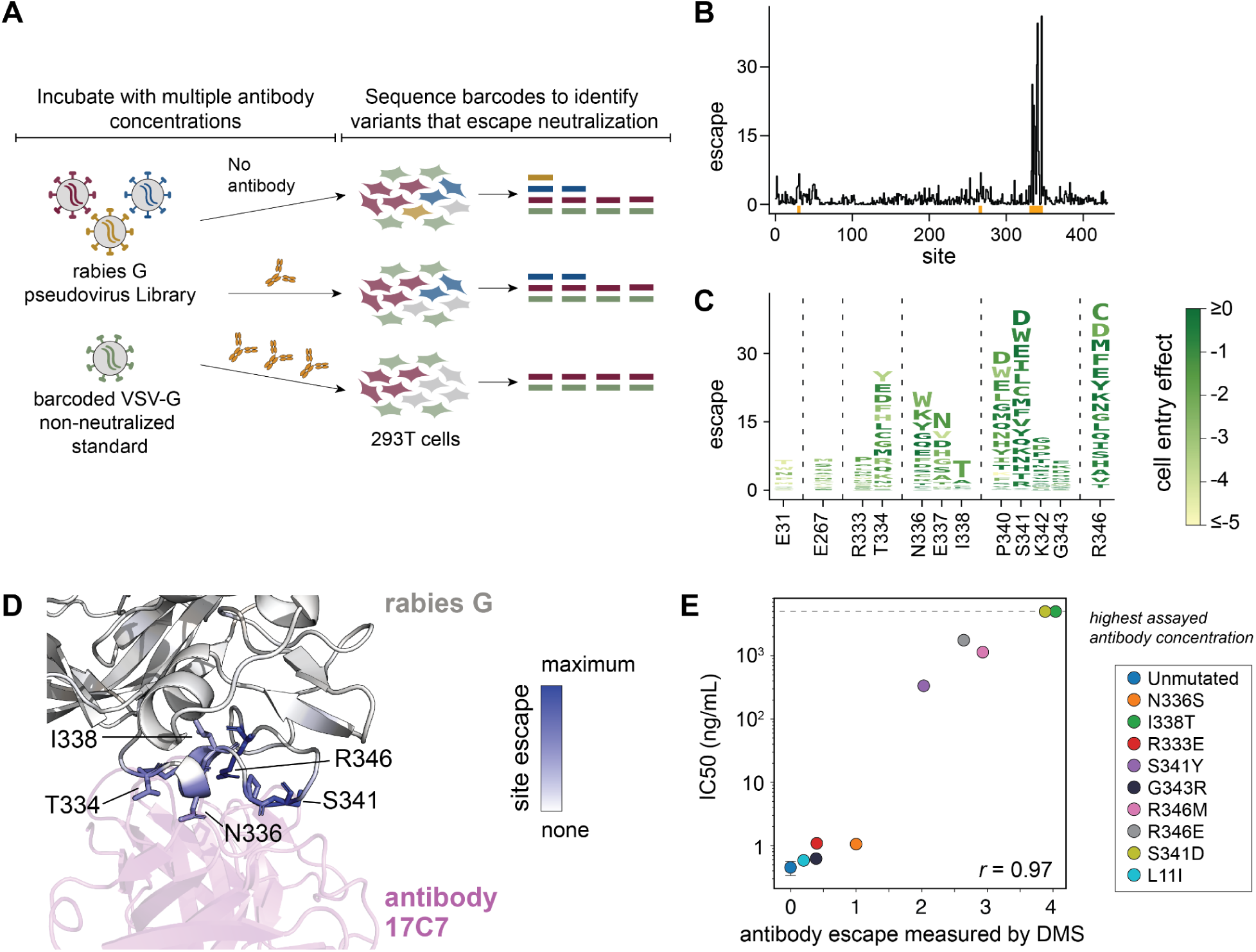
Deep mutational scanning of how G mutations affect antibody neutralization, as applied to antibody 17C7. (A) Workflow for mapping effects of mutations on antibody neutralization. The rabies G pseudovirus library is mixed with a “non-neutralized standard” consisting of barcoded VSV-G pseudovirus, then this pool is incubated with different concentrations of antibody and used to infect 293T cells. Barcodes are recovered from infected cells and deep sequenced to quantify the ability of each variant to infect cells at each antibody concentration. The VSV-G non-neutralized standard pseudovirus is not affected by anti-rabies G antibodies, and so is used to convert the sequencing counts into the fraction of each variant that is neutralized at each antibody concentration. (B) Line plot showing the total escape caused by mutations at each site in G for antibody 17C7. See https://dms-vep.org/RABV_Pasteur_G_DMS/escape.html for interactive visualizations of escape. Orange marks near the x-axis indicate sites zoomed in the logo plot in panel (C). (C) Logo plot with the height of each letter proportional to the escape caused by mutation to that amino acid for key sites of escape from antibody 17C7. The letters are colored according to the effect of each mutation on cell entry. (D) Structure of 17C7 and rabies G (PDB: 8A1E) with G colored by the total escape caused by mutations at each site (blue indicates sites where mutations cause the most escape). The antibody is shown in purple. (E) Validation assays comparing the escape measured in the deep mutational scanning (DMS) versus the IC50s measured in standard pseudovirus neutralization assays against antibody 17C7 for nine different single amino-acid mutants of G. Raw neutralization curves are in Fig S5. The mutations tested in these validation assays were chosen to span a range of escape values as measured in the deep mutational scanning. Pseudovirus with unmutated rabies G was measured in triplicate, and mean ± standard deviation is plotted.

We first applied this approach to monoclonal antibody 17C7, which is the active component of the Rabishield post-exposure prophylactic^16,33,46^ and has had its structure in complex with G determined by cryo-EM^12^. Our deep mutational scanning showed that mutations that escape neutralization by 17C7 occur predominantly at sites 334, 336-338, 340-342, and 346 (Fig 3B,C and interactive visualizations at https://dms-vep.org/RABV_Pasteur_G_DMS/escape.html). Many of these escape mutations are well tolerated with respect to G’s cell entry function (as shown by the color of letters in Fig 3C). The escape mutations all occur near the binding interface of 17C7 and G (Fig 3D). One of the escape mutations, I338T, does not directly contact the antibody but creates an N-linked glycosylation motif at N336 that is known to affect G’s antigenicity^47,48^. I338S does not appear as an escape mutation because, unlike I338T, it is highly deleterious for cell entry in the background of the Pasteur strain G (Fig 1C).

To validate the deep mutational scanning measurements of antibody escape, we created individual pseudoviruses encoding nine different mutations that had a range of effects on neutralization by antibody 17C7 in the deep mutational scanning. The effects of these mutations as measured in traditional neutralization assays performed with these pseudoviruses were highly correlated with the escape values measured in the deep mutational scanning (*r* = 0.97; Fig 3E, S5A), validating the accuracy of the approach.

### Comprehensive escape maps for eight monoclonal antibodies

We used deep mutational scanning to experimentally map all escape mutations for eight monoclonal antibodies targeting G (Table 1). These antibodies bind to different antigenic regions on G: two target antigenic region I (antibodies CR57^48,49^ and RVC20^34,50^), four target region III (17C7^16^, CR4098^49^, RVA122^25,34^, and RVC58^17,34^), and two bind outside previously described antigenic regions (CTB012^18^ and RVC68^34^). Antibody 17C7 has been clinically deployed in India since 2017^19,51^, and CTB012^18^ is one of the two active components of the SYN023 cocktail^52,53^ that recently concluded phase III clinical trials^54^. All eight antibodies neutralized pseudovirus with unmutated rabies G, albeit with varying potencies (Table 1, Fig S5B).

**Table 1.** Antibodies with escape mutations mapped in this study. IC50s are from pseudovirus neutralization assays (Fig S5B). Antigenic regions were taken from^34,73^ and antibodies marked “N/A” bind outside known antigenic regions. Pan-lyssavirus activity was defined based on strain panel testing from^34^.

| Antibody | Development Status | Antigenic Region | PDB ID | Pan-lyssavirus activity<sup>34</sup> | IC50 (ng/mL) |
| --- | --- | --- | --- | --- | --- |
| 17C7<sup>16</sup> | Clinically deployed in India as of 2017<sup>19,51</sup> | III<sup>34</sup> | 8A1E<sup>12</sup> | No | 0.35 |
| CR4098<sup>49</sup> | Part of CL184 cocktail<sup>49,80</sup>; development stopped after Phase II trial<sup>19,32</sup> | III<sup>81</sup> | N/A | No | 3.8 |
| RVA122<sup>34</sup> | N/A | III<sup>34</sup> | 7U9G<sup>25</sup> | Yes | 0.31 |
| RVC58<sup>34</sup> | Animal testing in cocktail with RVC20<sup>73</sup> | III<sup>34</sup> | N/A | Yes | 0.22 |
| RVC20<sup>34</sup> | Animal testing in cocktail with RVC58<sup>73</sup> | I<sup>34</sup> | 6TOU<sup>50</sup> | Yes | 0.36 |
| CR57<sup>82</sup> | Part of CL184 cocktail (development stopped after Phase II trial)<sup>19,32</sup>; part of cocktails in strain panel testing<sup>19,33</sup> | I<sup>82</sup> | 8R40<sup>61</sup> | Yes | 0.55 |
| CTB012<sup>18</sup> | Part of SYN023 cocktail; clinical trials ongoing<sup>52,53</sup> (Phase III complete<sup>54</sup>) | N/A<sup>18,35</sup> | N/A | Untested | 0.47 |
| RVC68<sup>34</sup> | N/A | N/A<sup>34</sup> | N/A | Yes | 7.6 |

The deep mutational scanning shows that each antibody is escaped by mutations at a relatively small number of sites that spatially cluster in the pre-fusion structure of G (Fig 4 and interactive visualizations at https://dms-vep.org/RABV_Pasteur_G_DMS/escape.html). For the four antibodies that have had their structures in complex with G experimentally determined, the major escape mutations all occur at sites in G near the antibody binding interface (Fig 5A,B). However, mutations at only a fraction of the contact sites strongly escape antibody neutralization (Fig 5A,B), a result consistent with prior work on antibody escape for other viruses^55–58^. In some cases, certain contact sites have no escape mutations simply because all mutations are too deleterious for G’s cell entry function (eg, site C228 for antibody RVC20 in Fig 5B), but in other cases contact sites tolerate many mutations but none cause appreciable escape from neutralization (eg, site E33 for antibody RVA122 in Fig 5A,B).

**Figure 4.**
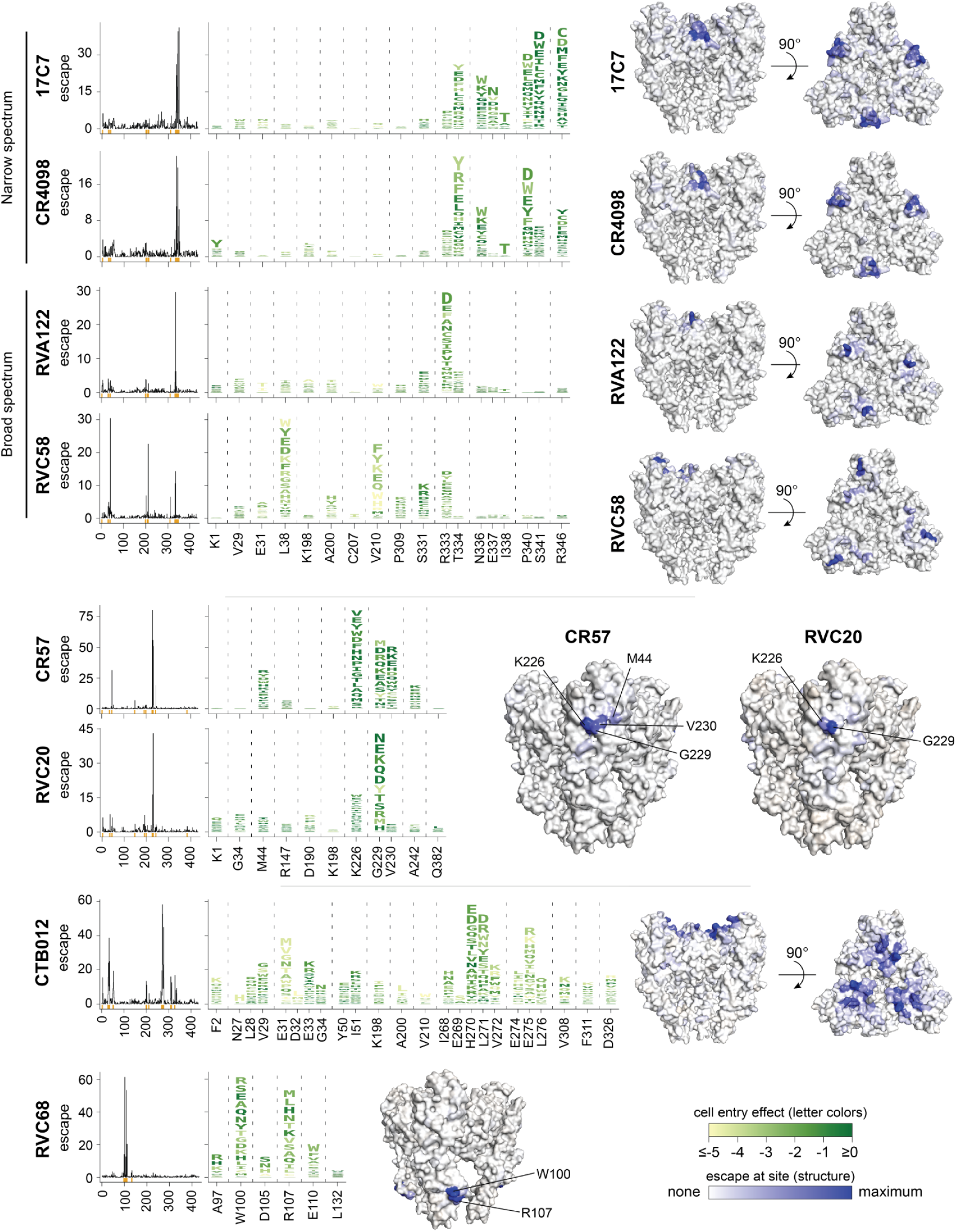
Comprehensive maps of escape mutations for eight monoclonal antibodies. For each antibody, the line plot shows the total escape caused by all mutations at that site; orange marks near the x-axis indicate sites zoomed in the logo plots. The logo plots show the escape caused by individual mutations at key sites, with letter heights proportional to escape value for mutation to that amino acid and letters colored according to the effects of mutations on cell entry (see color scale at the bottom right of the figure). The structures show pre-fusion rabies G (PDB: 7U9G) with residues colored by the total escape caused by mutations at each site (see color scale at bottom right of the figure). See Fig 5 for additional structural renderings of antibodies with experimentally solved structures in complex with G. See https://dms-vep.org/RABV_Pasteur_G_DMS/escape.html for interactive heatmaps and structure-based visualizations of escape.

**Figure 5.**
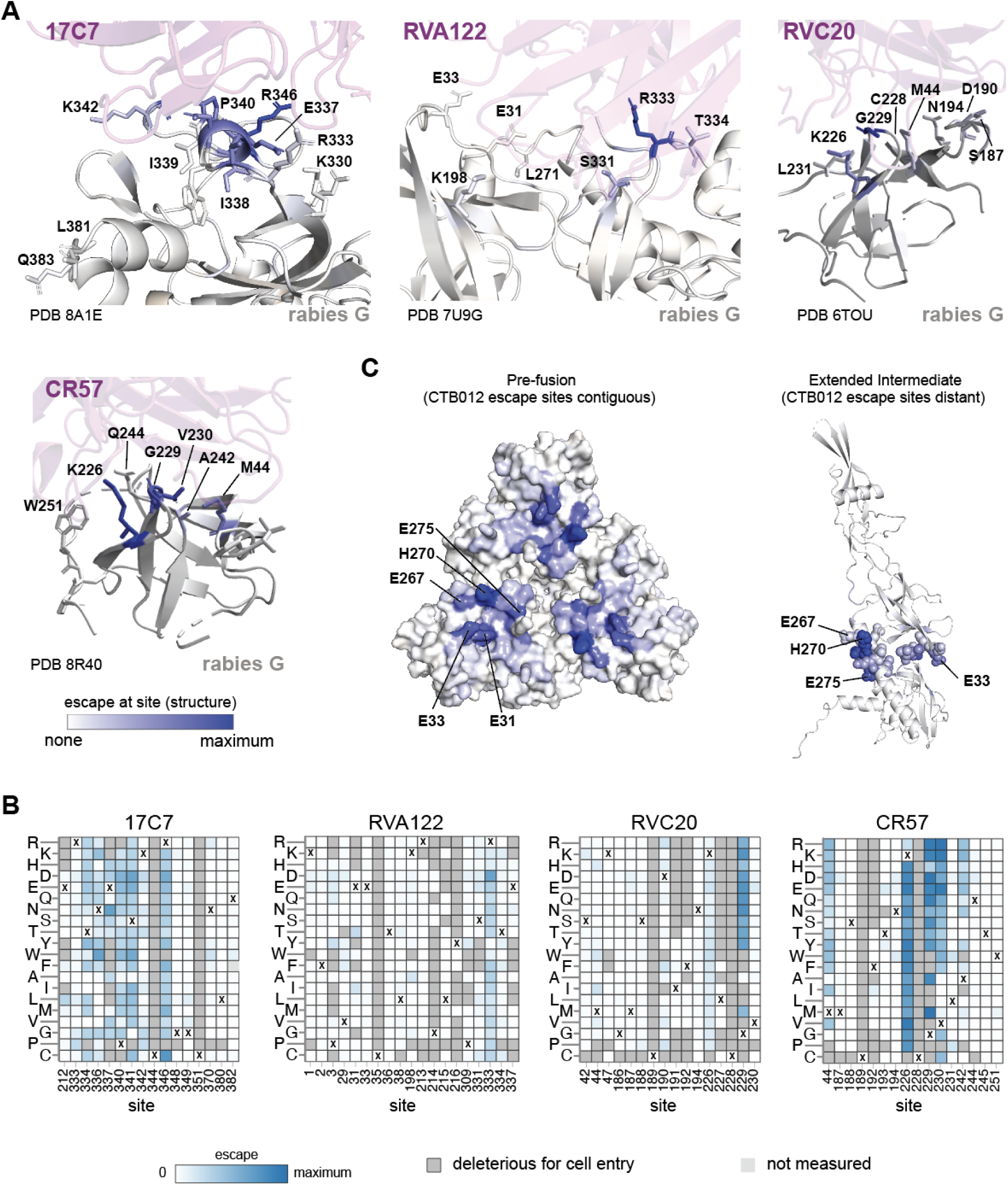
Mutations at only some of the antibody contact sites in G mediate escape. (A) G colored by extent of antibody escape at each site as measured in deep mutational scanning for the four antibodies with experimentally determined structures in complex with G (PDB 8A1E, 7U9G, 6TOU, and 8R40). The antibodies are shown in magenta. The color scale for G is defined at the lower left of the panel: white indicates no escape at the site, and blue indicates strong escape at the site. (B) Escape values for each mutation for all sites in G that contact each antibody in the experimentally determined structural complex. A site in G is defined as being in contact if any atom is within 4Å of the antibody. Mutations are colored white to blue based on the escape values measured in deep mutational scanning. Mutations that are too deleterious for cell entry to reliably measure their effect on antibody neutralization are shown in dark grey. The small number of mutations in light gray (eg. Q382F and K226H) have no measured escape due to insufficient coverage in the library. The amino-acid identity in the unmutated Pasteur G is indicated with a “X” for each site. (C) Structures of rabies G in pre-fusion (PDB 7U9G) and extended intermediate (PDB 6LGW) conformations colored with sites colored by the total escape from the CTB012 antibody. Escape sites are contiguous in pre-fusion rabies G but more distant in extended intermediate conformation. For visibility, key sites are shown in spheres in the extended intermediate. Color scale is the same as for (A).

Antibodies that target the same antigenic region of G can differ substantially in their escape mutations. For instance, 17C7, CR4098, RVA122, and RVC58 all bind to antigenic region III of G^34^. However, while 17C7 and CR4098 are escaped by mutations at very similar sets of sites, the sites of escape mutations for RVA122 and RVC58 are more distinct (Fig 4). Even when antibodies share sites of escape, the effects of individual mutations can differ. For instance, N336S^35^ and R346K^59,60^ escape 17C7 but not CR4098, although some other mutations at sites 336 and 346 escape both antibodies (Fig 4). Likewise, RVC20 and CR57 both bind to antigenic region I^34^ where they are believed to block the binding site for the alpha7 nicotinic acetylcholine receptor^22,61,62^ and prevent pH-driven conformational change^50,61^. Despite the similarity of their epitopes, the escape mutations for these two antibodies only partially overlap (Fig 4, 5A,B). For instance, both antibodies are affected by mutations to K226 and G229, but mutations at V230 and A242 only strongly escape CR57.

The escape maps also provide insight into why some antibodies have greater breadth against lyssaviruses than others. For instance, among the antibodies targeting antigenic region III, 17C7 and CR4098 only neutralize rabies whereas RVA122 and RVC58 also broadly neutralize non-rabies lyssaviruses (eg, Duvenhage lyssavirus, European bat 1 lyssavirus, and Australian bat lyssavirus)^34^. Unlike the narrow-spectrum antibodies (17C7 and CR4098), the broad-spectrum antibodies (RVA122 and RVC58) are escaped by mutations at fewer sites or sites with lower mutational tolerance. Specifically, RVA122 is mostly escaped by mutations at just one site: R333 (Fig 4, 5A,B), a site that is commonly mutated in vaccine strains^63–65^ but where mutations generally attenuate pathogenicity^66,67^. RVC58 is escaped by mutations at several sites, but many of those mutations, such as those at sites L38 and V210, are deleterious to G’s cell entry function (Fig 4, see color of letters in logo plots).

Two of the antibodies (CTB102 and RVC68) lack structures in complex with G and do not bind to previously defined antigenic regions, although the epitope of CTB102 has been partially defined by alanine mutagenesis and escape studies^18^. Our deep mutational scanning shows that CTB102 binds to sites on the apex of G that are spatially clustered on pre-fusion rabies G, primarily in the loops involving V29-E33 and I268-L275 (Fig 4, 5C). These loops undergo a substantial rearrangement in the extended intermediate conformation of G^24,27,31^, meaning that the epitope of CTB102 is only spatially contiguous in the pre-fusion conformation of G (Fig 5C). RVC68 has very broad neutralization activity across lyssaviruses^34^, but relatively poor neutralization potency against rabies G (Table 1). Our deep mutational scanning shows that RVC68 targets the base of the fusion domain of rabies G (Fig 4). The broad activity of this antibody against different lyssaviruses likely stems from conservation of the targeted sites in previously tested viruses^34^.

### Antibody-escape mutations are present in naturally occurring rabies strains

The G protein varies among natural rabies strains, and some strains have been identified that are resistant to specific antibodies, including clinically relevant antibodies^33,34,48,59^. Our deep mutational scanning enables systematic identification of antibody-resistant strains. We first examined the frequency of each amino-acid mutation among naturally occurring G sequences versus its antibody escape measured in our deep mutational scan (Fig 6A and interactive plot at https://dms-vep.org/RABV_Pasteur_G_DMS/natural_seqs.html). Mutations that cause at least some escape are present among natural G sequences for all antibodies, but the frequencies of these mutations and the magnitude of escape they cause differ markedly among antibodies. For example, many strong escape mutations from antibody 17C7 are at appreciable frequencies among natural sequences (Fig 6A). In contrast, antibodies such as RVC20 have only a few modest-effect escape mutations at appreciable frequency among natural sequences (Fig 6A).

**Figure 6.**
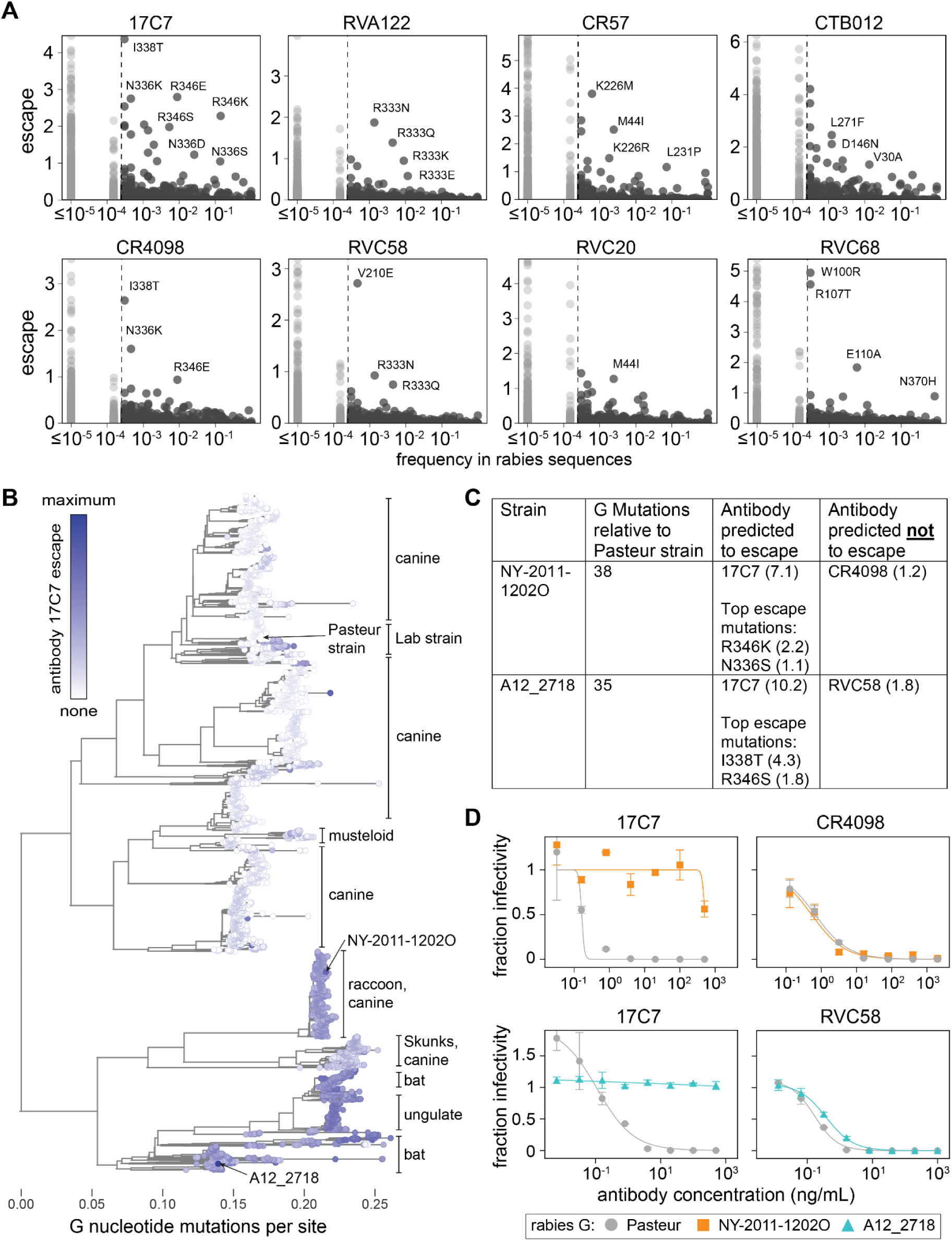
Naturally occurring antibody-escape mutations in rabies G. (A) Frequency of each G amino-acid mutation among naturally occurring rabies sequences versus the escape caused by that mutation in the deep mutational scanning of the Pasteur strain G. Each panel shows escape from a different antibody. Dashed vertical lines indicate mutations observed at least two times in natural strains; points in light gray indicate rare mutations. Mutations not observed in natural strains are assigned a frequency of 10^−5^ to enable plotting on a log scale. See https://dms-vep.org/RABV_Pasteur_G_DMS/natural_seqs.html for an interactive version of this plot which allows mouseover of individual points to see the mutation identities. (B) Phylogenetic tree of all publicly available 7,122 rabies G sequences colored by escape from antibody 17C7 as predicted by the summing the experimentally measured effects of all mutations relative to the Pasteur strain. White indicates no escape and blue indicates strong escape. See https://nextstrain.org/groups/jbloomlab/dms/rabies-G for interactive Nextstrain phylogenies showing the predicted escape of all strains for each antibody. The tree is rooted by Gannoruwa bat lyssavirus (Accession: NC_031988.1), a non-rabies lyssavirus also in Phylogroup I. Strains experimentally tested in validation neutralization assays are labeled. Common hosts for different clades are labeled. The interactive Nextstrain phylogenies provides the option to label all strains by host. (C) For validation neutralization assays, we identified antibodies predicted to be escaped or neutralized by two different natural strains. The table summarizes the strains and antibodies used in the validation assays. The numbers in parentheses after the antibody names and mutations in the last two columns are the escape values from the deep mutational scanning. (D) Neutralization curves of pseudovirus expressing G from the Pasteur, NY-2011-12012O, or A12_2718 strains against the indicated antibodies. Points indicate the mean ± standard error of technical duplicates.

We predicted each rabies strain’s escape from each antibody by summing the experimentally measured effects of all of its constituent G mutations relative to the Pasteur strain. We visualized these predictions via Nextstrain^68^ phylogenetic trees where sequences can be colored by their predicted escape (https://nextstrain.org/groups/jbloomlab/dms/rabies-G). As an example, a static view of the tree for all 7,122 publicly available G sequences colored by escape from antibody 17C7 is shown in Fig 6B. Although most strains lack escape mutations from 17C7 (white in Fig 6B), there are some strains predicted to strongly escape for this antibody (blue in Fig 6B). Most of these strains contain mutations at sites 336 or 346 (Fig S6A), both of which are sites where mutations are measured to strongly escape 17C7 in the deep mutational scan (Fig 4, 6A,C). Note that many of the sequenced strains that are predicted to most strongly escape 17C7 originate from infected bats or wild animals rather than dogs (Fig 6B), which are the most common source of human infections^69^.

As validation, we identified two natural strains predicted to strongly escape antibody 17C7, NY-2011-1202O and A12_2718 (Fig 6C). The G proteins of these two strains both differ from the Pasteur strain G by at least 35 amino-acid mutations. We generated pseudovirus expressing G from each of these strains and measured neutralization by 17C7 as well as another antibody predicted not to be escaped by each of the strains (Fig 6C,D). As predicted by the deep mutational scanning, both strains strongly escaped neutralization by 17C7 but retained neutralization by the control antibodies at levels similar to the Pasteur strain (Fig 6D).

To test how much the escape was explained by just the top escape mutations identified in the deep mutational scanning, we either removed each of the two top escape mutations (I338T or R346S) from the A12_2718 strain G or introduced them into the Pasteur strain G. Consistent with the deep mutational scan, removing the top escape mutation of I338T from the A12_2718 G partially restored neutralization by 17C7, while adding this single mutation to the Pasteur G made it fully resistant to neutralization by 17C7 (Fig S6B). Introducing the weaker escape mutation R346S to the Pasteur strain reduced but did not eliminate its neutralization by 17C7, also in agreement with the deep mutational scan (Fig S6B). While these measurements validate the ability of deep mutational scanning to predict escape of substantially divergent strains for the antibodies tested here, caution should always be exercised when extrapolating the deep mutational scanning to sequence-divergent strains. In some cases, epistasis may lead to shifts in the effects of mutations as observed in other viral proteins^70–72^.

## Discussion

Here, we used deep mutational scanning to measure how mutations to rabies G affect cell entry and antibody neutralization. The comprehensive measurements of how mutations affect G’s cell-entry function define which sites in the protein have the most capacity for evolutionary change, and shed light on how mutations affect G’s conformational dynamics in a way that could inform design of vaccine antigens. The measurements of how mutations affect antibody neutralization delineate key epitopes. This information can guide development of antibodies and antibody cocktails that are robust to G’s natural genetic diversity, as well as define key epitopes to preserve in vaccine design.

With respect to how mutations affect G’s cell-entry function, we find that some regions of G (eg, the base of the fusion domain and parts of the trimer apex) are intolerant of most mutations, but other regions can tolerate mutations without substantially impairing cell entry. In some instances, we can identify sites where mutational constraint appears to arise from protein contacts formed in just one of the pre-fusion or extended intermediate conformations. Mutations that differentially affect these two conformations could aid in the design of vaccine immunogens. The conformational dynamics of rabies G are hypothesized to hamper the neutralizing antibody response to vaccination by unmodified G, since some potently neutralizing antibodies target only pre-fusion G^13,25,26^. Several studies have therefore attempted to stabilize pre-fusion G by introducing proline mutations that block conformational changes^12,13^, similar to what has been done for other viral fusion proteins^42–44^. Our work suggests an additional mechanism of pre-fusion stabilization by identifying sites where mutations may destabilize the extended intermediate.

Our work better defines how neutralizing antibodies target G, and how these antibodies are affected by viral mutations. Delineation of which viral mutations escape anti-G antibodies is important since monoclonal antibodies are being developed for post-exposure prophylaxis, with some already in clinical use. We comprehensively mapped escape mutations for eight antibodies, including the antibody in Rabishield and several other antibodies under development for use against rabies^16–18,73^. Some of these antibodies have had their structures in complex with G previously determined, and our work shows that only some mutations at the antibody-G interface confer escape. Other antibodies that we mapped have not been structurally characterized, and for those antibodies our work defines their epitopes. Furthermore, our measurements help rationalize why antibodies differ in their breadth against lyssaviruses: the broadest antibodies tend to target sites in G where most mutations are detrimental to G’s cell-entry function.

Our deep mutational scanning can help identify which natural rabies strains are likely to escape specific antibodies. Strikingly, we find that escape mutations to all eight antibodies we examined are present among natural rabies strains—but escape mutations are far more common for some antibodies than others. We provide interactive phylogenetic trees that predict the escape of publicly available rabies G sequences based on the effects of their constituent mutations as measured in our experiments. These predictions are approximate because there can be epistasis among combinations of mutations^70–72^ (see the *Limitations of study* section below). However, we did experimentally validate several examples showing that our data can predict the susceptibility or resistance of natural strains to specific antibodies. Going forward, the deep mutational scanning data provide a powerful way to identify natural strains that are likely to harbor resistance mutations, although such predictions should always be independently validated given the potential for epistasis.

Overall, our work provides the first large-scale measurements of the functional and antigenic effects of mutations to rabies G. Our data can be interrogated in the context of G’s structure and evolution via interactive visualizations we have made publicly available (https://dms-vep.org/RABV_Pasteur_G_DMS/), which should facilitate their use in basic research, vaccine and antibody development, and surveillance of rabies virus evolution.

## Limitations of study

Our measurements were made using lentiviral particles pseudotyped with rabies G and infection of 293T cells. This pseudovirus system eliminates biosafety concerns associated with making mutants of actual pathogenic rabies viruses; however, it comes with the caveat that pseudovirus infection of 293T cells does not recapitulate all aspects of actual rabies virus infection *in vivo*^3,74,75^. Therefore, although our experiments do capture the core processes of G-mediated cell entry and antibody neutralization, they may not authentically reflect some more subtle aspects of real viral infection, such as differential usage of specific receptors^20,62,76^.

Our deep mutational scanning used G from the lab-adapted Pasteur vaccine strain, which has been passaged extensively in animals and cell lines^77^. Some of the mutations from this passaging enhance pseudovirus production^78,79^, but could alter some of G’s properties relative to natural strains. Additionally, there is always the potential for epistasis among mutations^70–72^, such that the effect of a mutation in the Pasteur G could differ for glycoproteins from other strains with multiple mutations. This limitation is especially relevant when using the deep mutational scanning to predict the antibody escape of other rabies strains (last section of *Results*). Although such predictions validated well in the examples we tested here, caution should always be exercised when predicting properties of divergent strains from measurements made on the Pasteur G.

Finally, like all high-throughput experiments, deep mutational scanning is subject to experimental noise. We performed all experiments in replicate using independently generated pseudovirus libraries, but in most of this paper we simply report the average of these replicates. For mutations of special interest, we recommend also examining the interactive heatmaps at https://dms-vep.org/RABV_Pasteur_G_DMS/ where you can mouseover points to look at the measurements for each individual replicate and view other quality control metrics (such as the number of unique variants in which each mutation was observed).

## Acknowledgments

We thank Brendan Larsen, William Hannon, and Sophie Shoemaker for technical advice and feedback. This work was supported by NIH grant R01 AI141707 to JDB. We thank the Genomics & Bioinformatics Shared Resource (RRID: SCR_022606) of the Fred Hutchinson Cancer Center (P30 CA015704). This research was supported by the Flow Cytometry Shared Resource, RRID:SCR_022613, of the Fred Hutch/University of Washington/Seattle Children’s Cancer Consortium (P30 CA015704). JDB is an Investigator of the Howard Hughes Medical Institute.

## Author contributions

Conceptualization: AKA, JDB; Methodology: AKA, CER, JDB; Software: CER, CRC, JDB; Investigation: AKA; Resources: NJ, NPK; Data curation: AKA, CER, CRC, JDB; Visualization: AKA, CRC, JDB; Writing–Original Draft: AKA, JDB; Writing–Review & Editing: AKA, CER, CRC, NJ, NPK, JDB; Supervision: JDB; Funding Acquisition: JDB.

## Supplemental Figures

**Figure S1.**
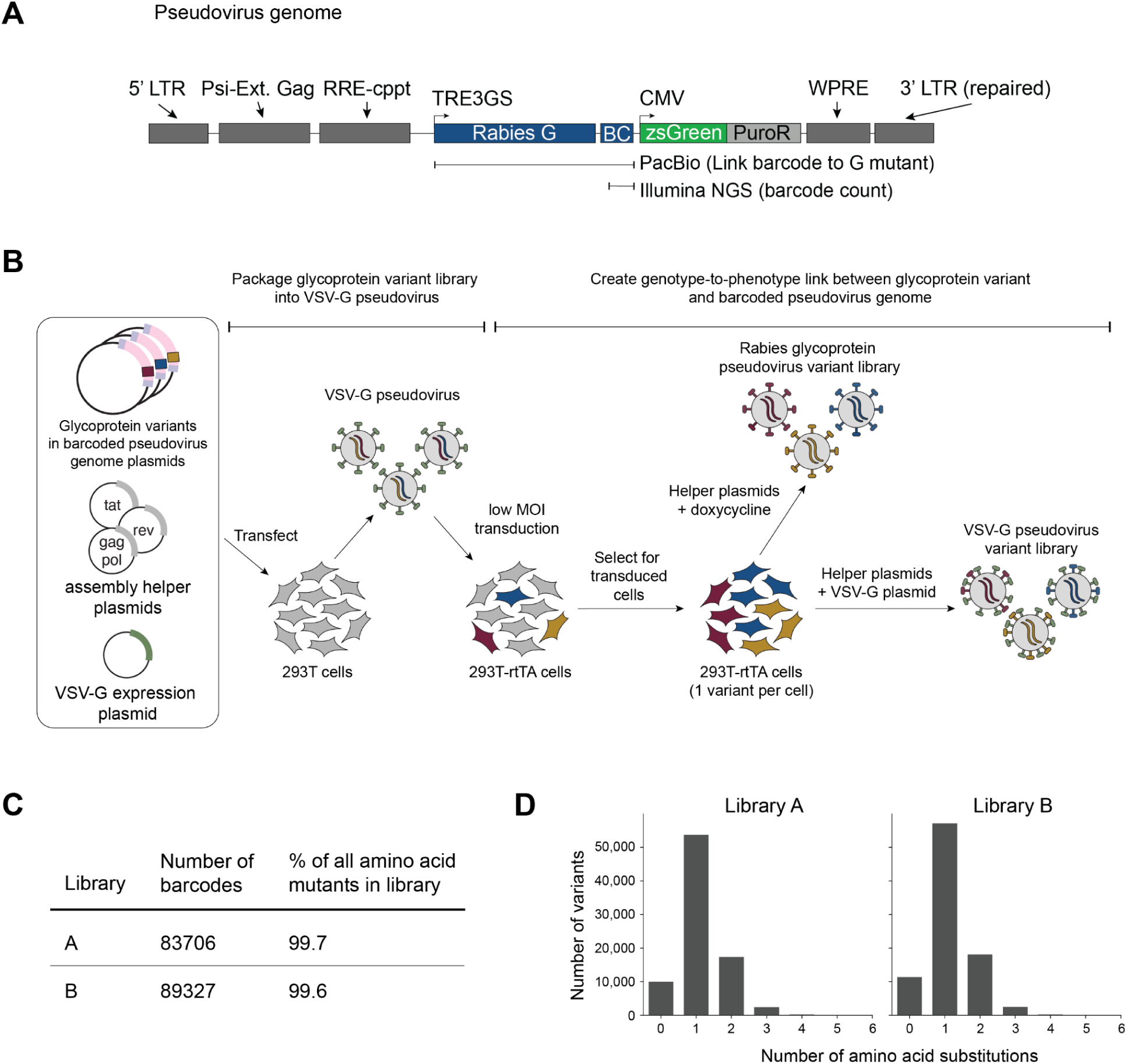
Pseudovirus deep mutational scanning of rabies G (Related to figure 1) (A) Schematic of lentiviral genome used to produce genotype-phenotype linked pseudovirus libraries. The genome encodes a dox-inducible rabies G mutant gene followed by a stop codon and 16 random nucleotides that function as a barcode. PacBio sequencing is used to link each barcode to a corresponding G mutant. In subsequent experiments, short-read sequencing of barcodes is used to measure cell entry for G mutants. There is a separate transcriptional cassette that constitutively expresses ZsGreen and a puromycin resistance gene. The 3’ LTR is repaired to enable reactivation of integrated proviruses. The pseudovirus genome employed in this study contains an extended Gag sequence (denoted as “Ext. Gag”), which may improve genome packaging into virions. (B) Workflow for generating genotype-to-phenotype linked pseudoviruses. We produce VSV-G pseudovirus particles with lentiviral genomes that encode the rabies G mutants and their barcodes. VSV-G pseudovirus are transduced into a 293T-rtTA expressing cell-line at a low MOI to ensure one genome integration per cell. Cells with integrated proviruses selected using puromycin. Library pseudoviruses are produced by transfecting only lentiviral helper plasmids (gag-pol, tat, and rev) and adding doxycycline to induce rabies G expression. VSV-G pseudovirus is simultaneously produced as a control to measure library composition for cell entry measurements by transfecting a VSV-G expression plasmid alongside the helper plasmids. (C) Number of barcoded variants and percentage of all 8,227 targeted amino acid mutations represented in each of the two replicate G pseudovirus libraries. (D) Distribution of number of amino acid mutations per G variant in each pseudovirus library. Most sequences contain only one mutation, but some contain no or multiple mutations.

**Figure S2.**
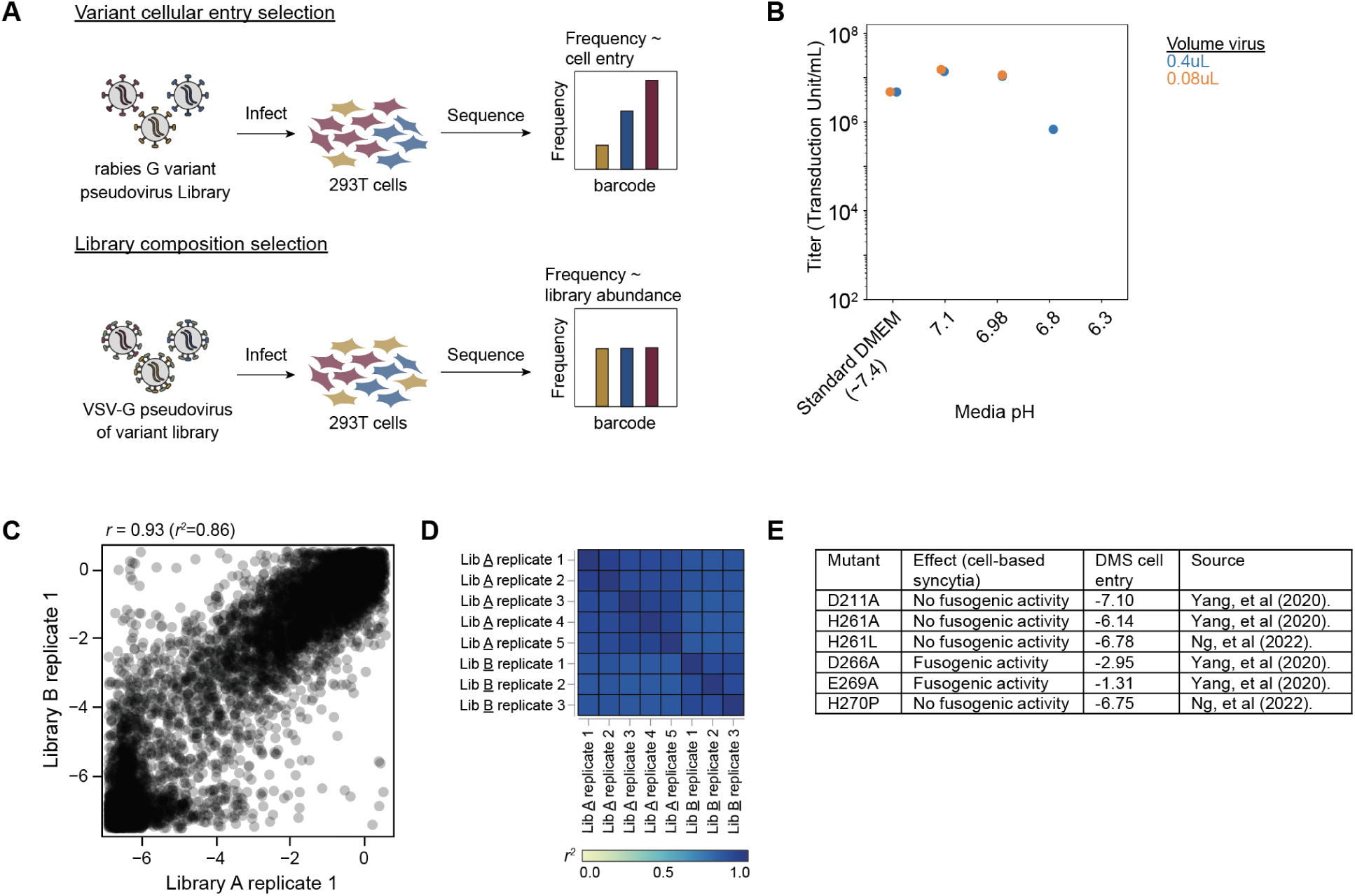
Measuring effects of rabies G mutations on cell entry (Related to Figure 1) (A) Workflow for measuring effects of mutations on cell entry. 293T cells are infected either with pseudovirus expressing only the rabies G mutants, or pseudovirus also expressing VSV-G. After infection, lentiviral genomes are isolated from infected cells and sequenced to identify the fraction of each variant that is able to infect cells. When the virions express only the rabies G mutant, only functional mutants will infect cells. But when VSV-G is also expressed, then all variants can infect cells. The cell entry score for each variant is quantified as the log of its frequency relative to unmutated rabies G in the condition expressing only rabies G versus the condition also expressing VSV-G. (B) The rabies G pseudovirus infections were performed using media with a pH adjusted to 7.1 as that yielded higher titers. Shown are the titers of a stock of pseudovirus expressing unmutated rabies G in media equilibrated to various pHs. (C) Correlation of the effects of G protein mutations on cell entry measured in a single experimental replicate with each of the two duplicate pseudovirus libraries. (D) Overall we performed five repeats of the cell entry measurements using library A and three repeats using library B. The heatmap shows the correlation of the mutation effects on cell entry between each pair of measurements. Throughout this paper, we use the median measurement across replicates for each mutation effect. (E) Summary of fusogenic activity for rabies G mutants assayed by cell-based syncytia assays as compared to deep mutational scan (DMS) cell entry scores. More negative deep mutational scanning scores correspond to reduced cell entry activity, with scores ≤ −5 indicating mutations that are as deleterious as most stop codons.

**Figure S3.**
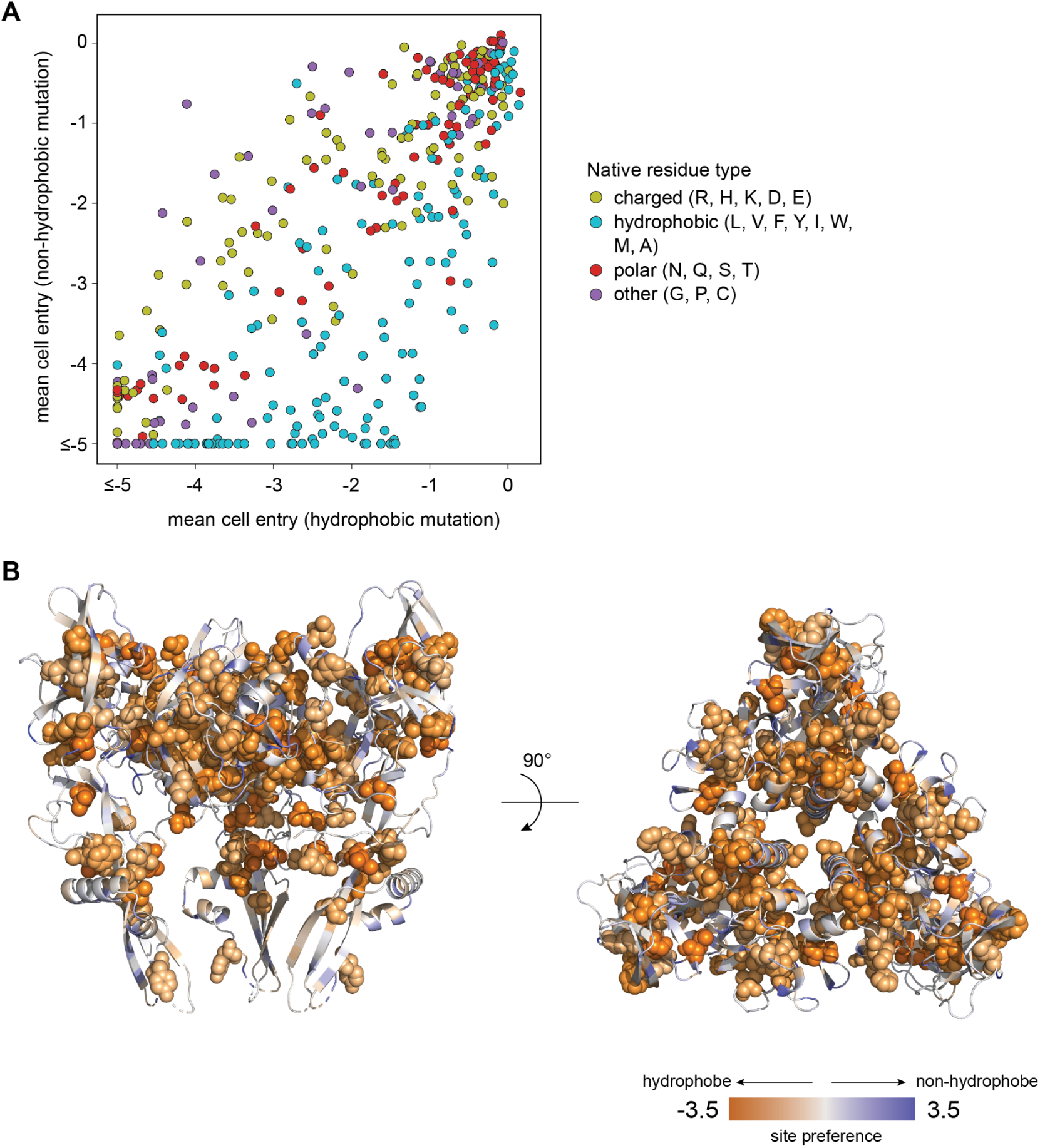
Constraint on hydrophobic residues in rabies G (Related to Figure 2). (A) Mean effect on cell entry for all mutations to non-hydrophobic amino acids versus the mean effect of all mutations to hydrophobic amino acids. Each point is a site, and is colored by whether the parental amino acid in G at that site is charged, hydrophobic, polar, or other as delineated in the key to the right of the plot. For the calculation of the mean effects, both charged and polar amino acids are grouped into the non-hydrophobic category, and the other category (G, P, and C) are excluded from the mean-effect calculations. (B) Pre-fusion rabies G structure colored by the preference of each site for hydrophobic versus non-hydrohobic amino acids. This preference score is calculated by subtracting the mean effect of all mutations to hydrophobic amino acids from the mean effect of all mutations to non-hydrophobic amino acids, so that negative values indicate a preference for hydrophobic residues. The structure is colored so that sites that prefer hydrophobic amino acids are colored orange, and ones that prefer non-hydrophobic amino acids are colored blue. The structure is shown as a cartoon with sites with a hydrophobicity preference more than one standard deviation below the mean of all sites shown in spheres. As expected, sites with hydrophobic preferences tend to be on the core rather than the surface of the protein.

**Figure S4.**
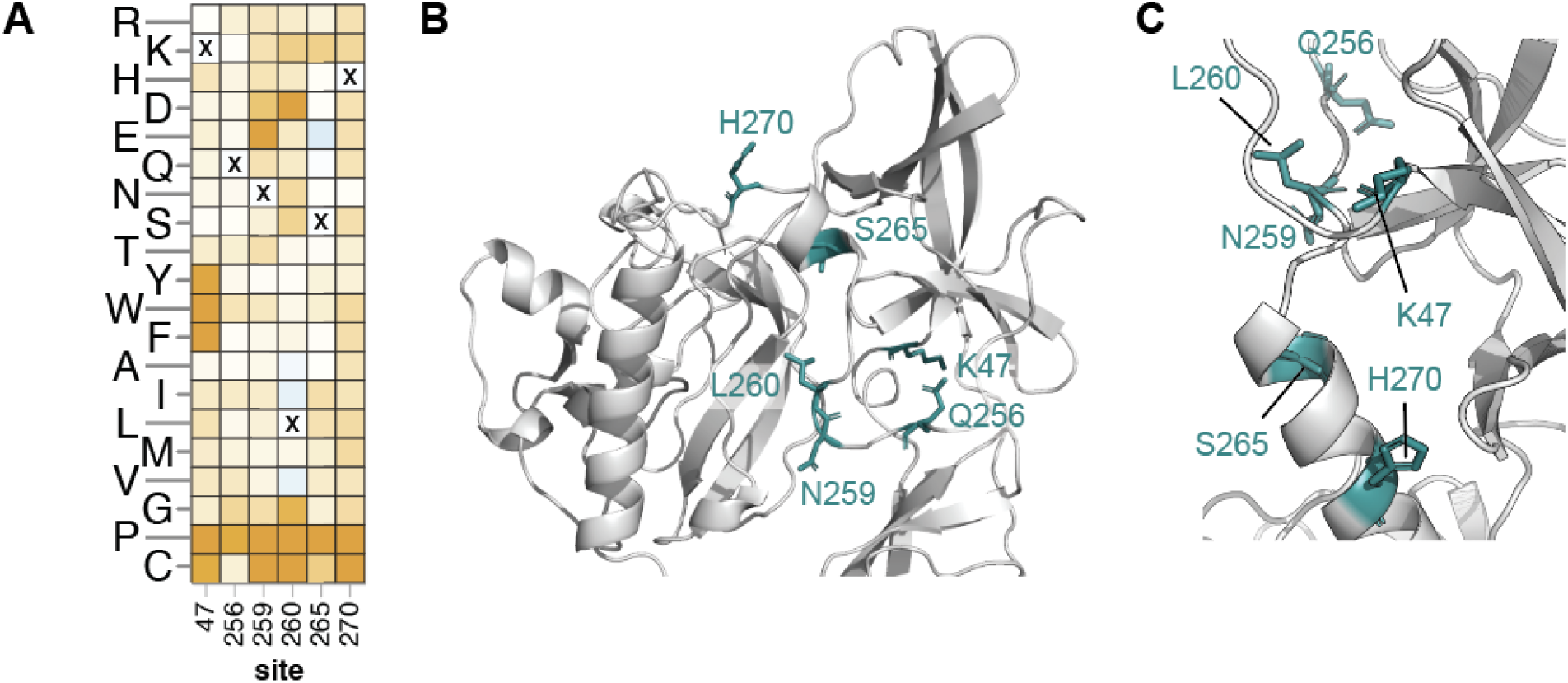
Sites where proline mutations are especially deleterious to cell entry (Related to Figure 2). (A) Effects of all mutations on cell entry (color scale as in Figure 1C) for a subset of sites where mutations to proline are highly deleterious but mutations to many other amino acids are well tolerated. (B) Zoomed in view of a single protomer of the pre-fusion trimer with the sites where proline mutations are highly deleterious shown in teal. (C) Zoomed in view of a single protomer in the extended intermediate conformation with the sites where proline mutations are highly deleterious shown in teal.

**Figure S5.**
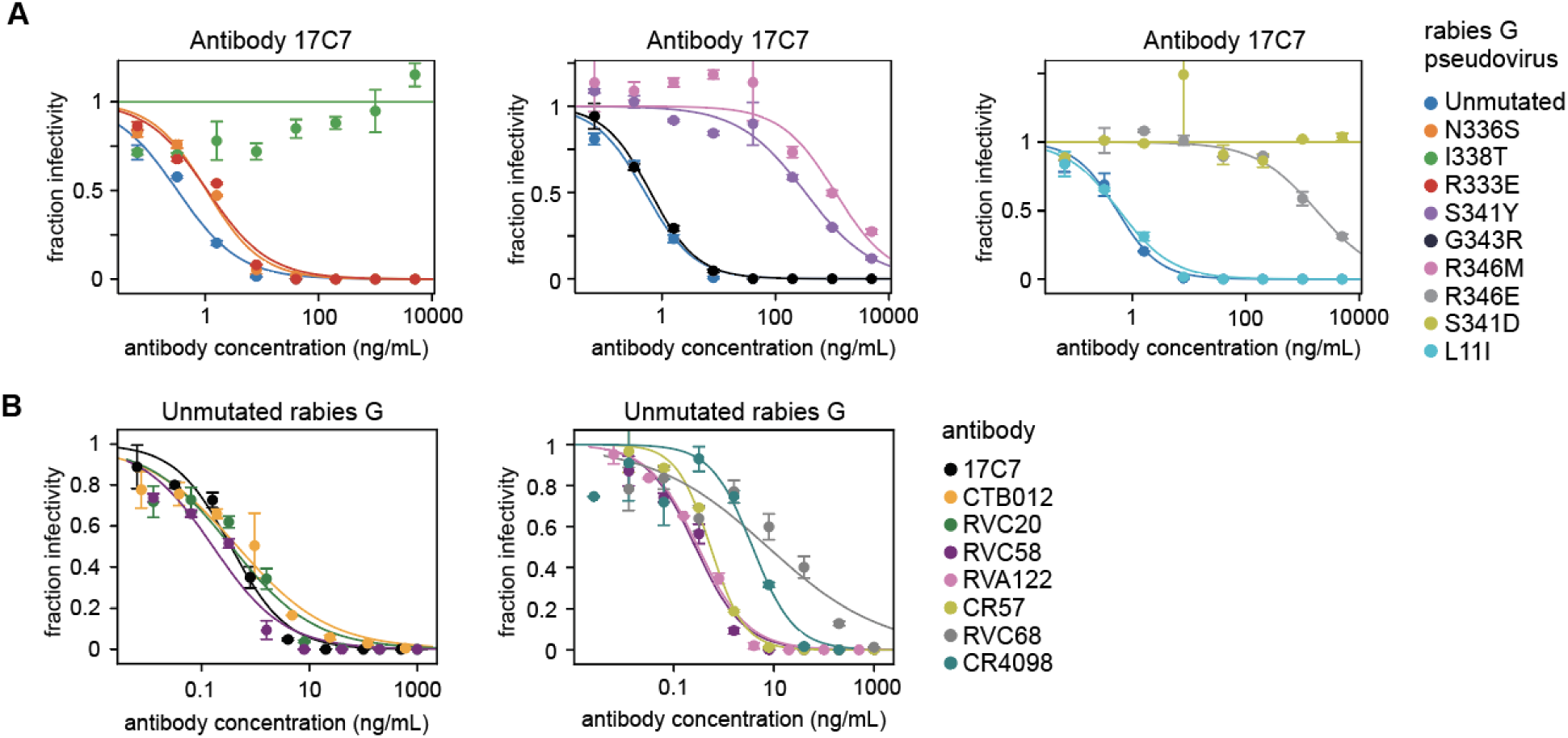
Neutralization curves (Related to Table 1 and Figures 3 and 4). (A) Neutralization curves for validation pseudovirus assays for nine rabies Pasteur strain G mutants against antibody 17C7. The neutralization assays for all nine mutants were split across three plates, each of which is shown in a separate panel. Unmutated rabies G pseudovirus was run on each plate as a control for plate-dependent variation. Each measurement point represents the mean ± standard error of two technical replicates. A Hill curve was fit to each mutant (solid line) to calculate the IC50. (B) Neutralization curves for unmutated Pasteur strain rabies G against eight antibodies studied. The neutralization assays were split across different experiments, each of which is shown in a separate panel. RVC58 was run on both plates as an internal control for variation.

**Figure S6.**
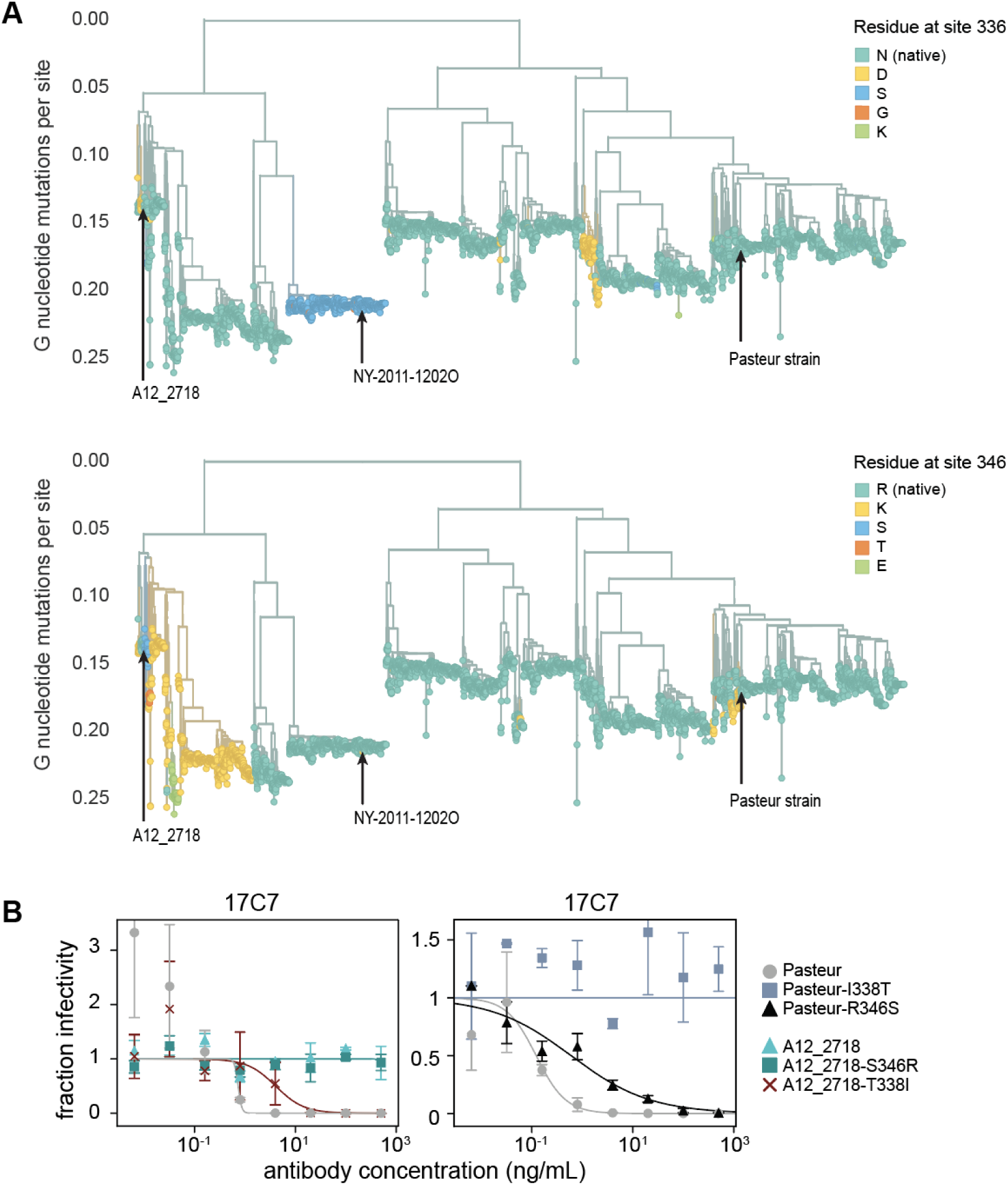
Escape mutations in circulating rabies G sequences (Related to Figure 6). (A) Phylogenetic trees of all publicly available rabies G sequences colored by the amino-acid identity at sites 336 and 346. Strains tested in the validation assays in Fig 5D are labeled. See https://nextstrain.org/groups/jbloomlab/dms/rabies-G for interactive trees that enable you to color the tree by the amino-acid identity at any site in G. (B) Neutralization curves of pseudovirus expressing single mutants in rabies G for both the Pasteur and A12_2718 strain backgrounds against antibody 17C7. Each panel represents assays run on a different plate, and points are the mean and standard error of technical duplicates. Note that as indicated in Fig 5C, I338T and R346S are the top two escape mutations found in strain A12_2718 relative to the Pasteur strain.

## Methods

### Data availability

See https://dms-vep.org/RABV_Pasteur_G_DMS/ for links to interactive visualizations of the data and CSVs containing the final inferred mutational results, including structure-based visualizations rendered with dms-viz^83^. See https://github.com/dms-vep/RABV_Pasteur_G_DMS for the full computer and all data starting with the barcode counts.

The interactive plots at https://dms-vep.org/RABV_Pasteur_G_DMS/ contain a number of options that enable different displays of the data or filtering by the confidence of the measurement. In the heatmaps you can mouseover points for details on the per-replicate measurements (note all mutations were measured in at least two libraries). There are also options to adjust the site summary statistic (eg, mean or sum of mutation effects at a site), zoom in on particular regions, and filter values as follows:

- minimum times_seen: average number of variants per-library that contain a specific mutation; mutations present in more variants are generally measured more accurately.
- maximum effect_std: maximum standard deviation of measurement across all replicates.
- minimum n_selections or minimum n_models: minimum number of replicates in which a particular mutation was measured.
- minimum max of effect at site or minimum max escape at site: only show sites for which at least one mutation has a measured value greater than this.
- minimum functional effect: for antibody-escape, gray out and ignore any mutations with an effect worse than this on cell entry; this filters highly deleterious mutations.

These filters are all set to reasonable values by default, but in some cases you may want to adjust them further. In the heatmaps, mutations that lack measurements that meet the above filtering criteria are shown in light gray; in the antibody-escape heatmaps mutations that are highly deleterious are shown in dark gray.

### Experimental model details

#### Rabies G used for deep mutational scanning

We performed the deep mutational scanning using a codon-optimized G protein from the Pasteur strain (NCBI reference number: NC_001542.1, locus number: NP_056796), which is a vaccine strain^15^ with a G protein that has had its structure determined by cryo-EM^12,25^. See https://github.com/dms-vep/RABV_Pasteur_G_DMS/blob/main/Additional_Data/LibrarySpecs/Rabies-G_Pasteur-fullORF.fasta for the full length codon-optimized G used in this study. To maintain consistency with the standard numbering convention for G^19,25,50^, we assign the number 1 to the first protein site in the ectodomain (site 20 in the full-length protein encoded by the G ORF). Note that the interactive Nextstrain trees at https://nextstrain.org/groups/jbloomlab/dms/rabies-G have options to number sites either using this standard G ectodomain numbering sequence or starting with the N-terminal methionine of the full-length protein.

#### Cells

As target cells for infections to measure mutation effects on cell entry and antibody escape, we used 293T cells from ATCC (CRL-3216). For producing pseudovirus libraries from integrated cells (Fig S1), we used a previously described^36^ 293T-rtTA cell line. We cultured all cell lines in D10 media (Dulbecco’s Modified Eagle Medium supplemented with 10% heat-inactivated fetal bovine serum, 2 mM l-glutamine, 100 U/mL penicillin, and 100 mg/mL streptomycin). When working with the 293T-rtTA cell line, we used tetracycline-negative fetal bovine serum (Gemini Bio, Ref. No. 100-800). For generating pseudovirus libraries that either express rabies G mutants or VSV-G, we used Dulbecco’s Modified Eagle Medium without phenol red as we found this was helpful for pseudovirus concentration as it avoided also concentrating phenol red.

We noted that titers for pseudovirus expressing rabies G were higher at a pH of 7.1 (Fig S2B). For all infections with the rabies G pseudovirus libraries and single mutant validation assays, we formulated cell culture media that adopts a pH of 7.1 at 37°C and 5% CO_2_ based on a prior published protocol^39^, supplemented with 4 µg/mL polybrene (EMD Millipore, Ref. No. TR-1003-G). We used powdered Dulbecco’s Modified Eagle Medium (Corning™ 90013PB) formulated without L-glutamine, pyruvate, sodium bicarbonate, and phenol red. This media base was reconstituted in Milli-Q water per manufacturer’s instructions and supplemented with 10% tetracycline-negative heat-inactivated fetal bovine serum, 2 mM l-glutamine, 100 U/mL penicillin, 100 mg/mL streptomycin, and sodium bicarbonate to a final concentration of ∼0.75 g/L. Prior to use, we confirmed the pH of media after incubation for 18 hours 37°C and 5% CO_2_.

#### Antibodies

Antibodies were produced by Genscript as human IgG1 kappa isotypes. The sequences for heavy and light chains are at (https://github.com/dms-vep/RABV_Pasteur_G_DMS/blob/main/Additional_Data/AntibodySequences/RABV-G_DMS_AntibodySequences.fasta). Sequences were obtained from original patents or publications. RVA122, RVC58 and RVC58 originated from patent WO2016/078761A1^84^, CR4098 and CR57 originated from patent US7959922B2^85^, RVC20 came from original publication^50^, and CTB012 from patent WO2013174003A1^86^.

### Method details

#### G plasmid mutant libraries

The lentiviral backbone is schematized in Fig S1A and is mostly identical to the backbone previously used for pseudovirus deep mutational scanning of other viral entry proteins^36,37,87–90^. However, to enhance packaging of genomes in pseudovirus particles, we modified the lentiviral backbone to contain an extended Gag sequence^91^. A full plasmid map that contains the lentiviral backbone encoding the codon-optimized rabies G sequence is at (https://github.com/dms-vep/RABV_Pasteur_G_DMS/blob/main/Additional_Data/PlasmidSequences/3969_V5LP_pH2rU3_ForInd_ExtGag_Humes_RABV-G_Pasteur-Genscript3_CMV_zsGT2APurR.gb).

We mutagenized G starting two residues prior to the beginning of the ectodomain and continuing up to site 431 (so sites −2 to 431 in the standard numbering scheme). We designed the mutant library to encode all 19 mutant amino-acids at each site, and also to include 20 stop codons at alternating positions from the start of the mutagenized region as negative controls for cell entry. We ordered the mutant library as a site-saturation variant library from Twist Biosciences. The final Twist quality control report for the library is available at (https://github.com/dms-vep/RABV_Pasteur_G_DMS/blob/main/Additional_Data/LibrarySpecs/Q-293667_Final_QC_Report%20(9).xlsx). We noted that 38 mutants from the overall library were missing. Only site H384 (site 403 in Quality Control report) failed in synthesis, with 14 amino acid mutations missing.

We appended barcodes consisting of 16 random nucleotides to the mutagenized rabies G gene fragments downstream of the stop codon using PCR largely as described previously^87^. To generate biological replicate libraries, Library A and Library B, we performed two separate barcoding PCR reactions. For each barcoding PCR, we combined 5 ng of the library template from Twist (1 µL), 1.5 µL of forward primer (BC_FOR, 10 µM stock), and 1.5 µL of reverse primer (BC_REV, 10 µM stock), 21 µL of molecular biology grade water with KOD Hot Start Master Mix (ThermoFisher, Ref. No. 71842-4). See https://github.com/dms-vep/RABV_Pasteur_G_DMS/blob/main/Additional_Data/PrimerSequences/RABV-G_Sequencing-Primers.fasta primer sequences. PCR cycling conditions were as follows:

1. 95°C, 2 min
2. 95°C, 20 sec
3. 55.5°C, 20 sec, cooling at 0.5°C/sec
4. 70°C, 1 min
5. Return to Step 2, 9 cycles
6. 12°C hold

The PCR product was run on a 0.8% agarose gel and a fragment of the correct size was excised for clean up using a NucleoSpin Gel and PCR Clean-up kit (Macherey-Nagel, Ref. No. 740609.5) followed by Ampure XP beads (Beckman Coulter, Ref. No. A63881) and elution in water.

To prepare the recipient lentiviral vector, we digested a lentiviral plasmid that contains mCherry in the location where the G viral entry protein is cloned (4016_V5LP_pHrU3_ForInd-Extgag_mcherry) with XbaI and MluI-HF (NEB, Cat. No. R0145S and R3198S respectively). The digested product was gel purified and cleaned with Ampure XP beads as for the PCR product.

For each library, barcoded mutants were cloned into the digested lentiviral vector using HiFi assembly at a 2:1 insert to vector molar ratio, at 50°C for 1 hour. HiFi products for each library were cleaned with Ampure XP beads and transformed into 10-beta electrocompetent cells (NEB, Ref. No. C3020K) with a MicroPulser Electroporator (BioRad, Ref. No. 1652100), shocking at 2 kV for 5 milliseconds. Transformants were plated onto LB+ampicillin plates overnight at 37°C. To ensure the library was not bottlenecked and that the barcodes that are ultimately assigned to different variants are unique, we sought to maximize the number of colonies for each library. (Note that the library size is determined at the step of creating the cell-integrated libraries as schematized in Fig S1b, not at the plasmid creation stage; the plasmid library size must substantially exceed the final desired size of the cell-integrated libraries as there can be recombination during production of the lentiviruses that are integrated into the 293T-rtTA cells.) Library A contained 9.2 ✕ 10^6^ colonies, and Library B contained 7.8 ✕ 10^6^ colonies. Colonies were scraped off of plates into liquid luria broth, and plasmids were maxi-prepped with a HiSpeed Plasmid Maxi Kit (Qiagen, Ref. No. 12662).

#### Cell-stored rabies G mutant library

A deep mutational scan requires the ability to link the rabies G mutant expressed on each pseudoviruses’s surface with the barcode encoded in the genome. To accomplish this, we used the previously described^36,37,87,88,90^ approach shown in Fig S1B, which involves creating a library of cells each containing a single integrated lentiviral genome encoding a different barcoded G mutant. As schematized in Figure S1B, we first generated VSV-G expressing pseudoviruses encoding the rabies G mutants in their genomes. Using VSV-G as the viral glycoprotein ensures that cell entry occurs independently of the function of the rabies G encoded in the lentiviral genome. Next we used these VSV-G pseudovirions to transduce the 293T-rtTA cell line at a low multiplicity of infection (<0.01) to ensure single genome integration into these cells, followed by selection using puromycin (Sigma, Ref. No. P8833-25MG).

Specifically we produced VSV-G pseudovirus that package the rabies G encoding genomes by transfection at the scale of 30 mL. Approximately 4 ✕ 10^6^ cells were plated in each of three 10 cm dishes. Each plate was transfected with 5µg of the plasmid encoding the lentiviral backbone with barcoded G mutants, 1.25µg each of plasmids 26_HDM-Hgpm2 (Gagpol), 27_HDM-tat1b (Tat), 28_pRC-CMV-Rev1b (Rev), and 29_HDM-VSV-G (these plasmids are all available on AddGene under IDs 204152, 204154, 204153, and 204156; see https://www.addgene.org/browse/article/28237987/). Plasmids were transfected using the BioT transfection reagent (Bioland Scientific, Ref. No. B01-02) per manufacturer’s instructions. After 48 hours, supernatant from the plates was collected and filtered using a 0.45 µm syringe filter (Corning, Ref. No. 431220), and this supernatant was stored at −80°C until further use. To measure transducing units/mL (TU/mL) of this virus, we thawed an aliquot and transduced 293T cells and measured counts of ZsGreen-positive cells by flow cytometry.

As there were ∼8,000 amino acid mutations in the rabies G library, we aimed for 80,000 variants so that each mutant would have approximately 10 barcodes. We followed a prior protocol^37^ as described to transduce 293T-rtTA cells and collect transduced cells. Briefly, 6-well plates were seeded with 600,000 293T-rtTA cells. At the time of infection, we counted cells in a subset of wells and calculated transducing units of VSV-G pseudovirus to achieve a multiplicity of infection of 0.01. We transduced approximately 25 wells of cells for each library. After 48 hours, we measured transduction for some wells by flow cytometry counting ZsGreen cells and observed a final multiplicity of infection of 0.006. Using this percentage and the cell counts at the time of infection, we back-calculated the number of initial infected cells per well and pooled cells from wells to achieve approximately 80,000 infected cells for each library. Note that this entire process was performed separately for each of the two duplicate barcoded plasmid libraries described above.

Each library was pooled into 50 mL aliquots in tetracycline-negative D10 media supplemented with 0.75 µg/mL puromycin to select for transduced cells, which constitutively express the puromycin resistance gene in the lentiviral genome. These cells were initially plated in two 15 cm plates for each library. Cells were passaged and expanded in tetracycline-negative media with 0.75 µg/mL puromycin until only ZsGreen-positive cells were observed by microscopy. To expand the cell libraries, each was expanded into four 5-layer flasks (Falcon, Ref. No. 353144) in the absence of puromycin. To prevent future bottlenecking, we passaged cells at numbers in vast excess of the variants (eg. minimum 20 ✕ 10^6^ cells). Cell libraries were stored in aliquots of over 20 ✕ 10^6^ cells per aliquot in the vapor phase of liquid nitrogen. To protect the cell libraries during the freeze-thaw process, we froze cells in freezing media, composed of tetracycline-negative D10 media supplemented with 10% DMSO (by volume) and an additional 10% (by volume) of tetracycline-negative fetal bovine serum.

#### Production of rabies G and VSV-G expressing pseudovirus libraries

After creating the library of cells each of which encodes only one barcoded rabies G mutant lentiviral backbone, we produced pseudovirus particles expressing either rabies G or VSV-G on their surface (Fig S1B). The VSV-G expressing pseudoviruses were used for sequencing to link the barcode to the rabies G mutants (see below) and to control for variation in library composition during experiments to measure the effects of mutations on cell entry.

To produce rabies G the pseudovirus library, we seeded a minimum of three 5-layer flasks with ∼150 million cells each in phenol-free, tetracyline-negative D10 media. To each flask, doxycycline was added to a final concentration of 1 µg/mL to begin inducing expression of rabies G mutants from integrated genomes. At the time of transfecting helper plasmids 24 hours later, the D10 media was swapped for ∼150mL of fresh media supplemented with 1 µg/mL of doxycycline. For each flask, 60µg each of 26_HDM-Hgpm2 (Gagpol), 27_HDM-tat1b (Tat), and 28_pRC-CMV-Rev1b (Rev) expression plasmids were combined with 9.2mL of DMEM and 276µL of BioT transfection reagent. This transfection mix was incubated at room temperature for 15 minutes then added to the 5-layer flasks. Approximately 40 hours later, media supernatant was collected and sterile filtered using 0.45µm SFCA 500mL Rapid-Flow filter units (Nalgene, Ref. No. 09-740-44B). Filtered supernatant was concentrated ∼10-fold using Pierce™ PES, 100K MWCO, 20–100 mL Protein Concentrators (Thermo, Ref. No. 88537) by centrifugation at ∼1210*g* for 30-40 minutes, which was sufficient to achieve pseudovirus library titers of 10^6^ transduction units/mL. Until further use, the virus was aliquoted and stored at −80°C.

We produced VSV-G expressing pseudovirus by seeding two 15 cm plates each with 20 million library cells at the same time of seeding 5-layer flasks. To reduce expression of rabies G in these cells, we seeded cells in tetracycline-negative media and maintained cells in tetracycline-negative media when producing pseudovirus (note some rabies G will still be expressed off the lentiviral Tat-driven promoter). As with rabies G library pseudovirus, we replaced media immediately prior to transfection. To each plate, we transfected 9µg each of HDM_Hgpm2 (Gagpol), 27_HDM_tat1b (Tat), and pRC_CMV_Rev1b (Rev) expression plasmids with 3.75µg of 29_HDM_VSV-G expression plasmids. These plasmids were combined with 1.5mL of DMEM and 45µL of BioT. After incubation for 15 minutes, this transfection was added drop-wise to cells as per manufacturer instructions. Forty hours later, supernatant was sterile filtered using a 0.45 µm syringe filter (Corning, Ref. No. 431220) and aliquoted for storage at −80°C.

To determine the necessary volume of virus for experiments, we measured titers in transducing units/mL (TU/mL). We note that measuring titers may underestimate the viral particles used in all assays. Subsequent experiments rely on mini-prepping unintegrated lentiviral genomes, which are in excess of integrated genomes measured by titers^92–94^. However, we utilized titration measurements to avoid the risk of bottlenecking the library during infection experiments.

For viral titer measurements, small aliquots of virus were thawed at room temperature. To determine the TU/ml, we made a 5-fold dilution series of virus and measured the percentage of 293T cells infected at multiple concentrations of virus using flow cytometry for the ZsGreen encoded in the lentiviral genome at 40-48 hours post-infection. For VSV-G library pseudovirus, we performed these infections in standard tetracycline-free D10 media (pH of 7.4). For rabies G pseudovirus, we diluted virus in pH 7.1 media and swapped cell media for pH 7.1 media because (as described above) low pH media yielded better titers for rabies G. Prior to combining rabies G library pseudovirus and cells, media was equilibrated to the correct pH at 37°C and 5% CO_2_ in a tissue culture incubator for 1 hour. For antibody escape infection assays described later, we also measured barcoded VSV-G mCherry pseudovirus in these low pH conditions to plan the volume required to achieve 1-2% of titers relative to rabies G pseudovirus.

#### Long-read sequencing to link barcodes with rabies G mutants

We used PacBio long-read sequencing to link barcodes with the encoded rabies G variants in the lentiviral genome, largely as described previously^36,37,87,88^. Because the pseudo-diploid nature of HIV-derived lentiviruses leads to recombination during the infection of 293T-rtTA cells for generating single integrants, we performed the PacBio sequencing on pseudoviruses generated from the integrated cells rather than the plasmids initially used to generate the pseudovirus to create these cells^36^.

Specifically, 24 hours prior to infections, we plated ∼500,000-600,000 293T cells per well in poly-L-lysine coated 6-well plates. We aimed to transduce cells with ∼12 million transducing units of VSV-G expressing pseudovirus for each library. VSV-G expressing pseudovirus for each library was separately concentrated by centrifugation using Amicon Ultra 15mL/100kD MWCO Centrifugal filters (EMD Millipore, Ref. No. UFC9100024). These concentrated viruses were resuspended to a volume of ∼7mL of D10 media. We removed media from 3 wells of cells and added 2mL of virus to each well. After infection for 12-14 hours, each well of cells was separately miniprepped using a QIAprep Spin Miniprep Kit (Qiagen, Ref. No. 2710) to recover non-integrated reverse-transcribed lentiviral DNA. To improve yield, we heated miniprep columns during the elution step to 55°C for ∼5 minutes prior to centrifugation. After elution, we pooled the miniprep eluent each library for subsequent PCR steps.

We performed a two-step PCR product to generate amplicons for long-read sequencing as originally described in Dadonaite et al^36^. To detect strand exchange events originating from the PCR, we performed a first round of PCR in two separate reactions that introduces a single nucleotide tag (either a G or C) at the 5’ or 3’ ends of the amplicon. For each reaction, we combined 20µL of KOD Hot Start Master Mix, 15µL of miniprepped DNA, and molecular biology grade water. For primers, we either added 1µL each of 5_PacBio_primer_G and 3_PacBio_primer_C (both at 10µM stock) or 1µL each of 5_PacBio_primer_C and 3_PacBio_primer_G (both at 10µM stock). See https://github.com/dms-vep/RABV_Pasteur_G_DMS/blob/main/Additional_Data/PrimerSequences/RABV-G_Sequencing-Primers.fasta for PacBio Round 1 primers. PCR cycling conditions were as follows:

7. 95°C, 2 min
8. 95°C, 20 sec
9. 60°C, 10 s, cooling at 0.5°C/sec
10. 70°C, 1 min
11. Return to Step 2, 7 cycles
12. 70°C, 1 min
13. 4°C hold

The PCR products were cleaned with 50µL of Ampure XP beads (1:1 volume ratio with PCR reaction) and eluted in 35µL of elution buffer. For each library, we pooled equal volumes of each separate PCR reaction into a template master mix for the second step of the PCR. For each step 2 PCR reaction, we combined 21µL of the round 1 product, with 2uL each of 5_PacBio_Rnd2 (10µM stock) and 3_PacBio_Rnd2 (10µM stock) primers. See https://github.com/dms-vep/RABV_Pasteur_G_DMS/blob/main/Additional_Data/PrimerSequences/RABV-G_Sequencing-Primers.fasta for the PacBio Round 2 primer sequences. PCR cycling conditions were as follows:

14. 95°C, 2 min
15. 95°C, 20 sec
16. 70°C, 1 sec
17. 60°C, 10 s, cooling at 0.5°C/sec
18. 70°C, 1 min
19. Return to Step 2, 10 cycles
20. 70°C, 1 min
21. 4°C hold

Each PCR reaction was cleaned with 50µL of Ampure XP beads (1:1 volume ratio with PCR reaction) and eluted in 40µL of elution buffer. We combined PCR reactions for each library together and verified by TapeStation that the amplicons were monodisperse and of correct length prior to sequencing. We sequenced each library on a single SMRTcell with a movie length of 30 hours on a PacBio Sequel IIe sequencer. To sequence to a higher depth, we later sequenced Library B on an additional SMRTcell and appended this to our analysis. Computational analysis of long-read sequencing to establish the barcode and variant linkages is detailed in the “PacBio sequencing analysis” section. That analysis checked for swapping of the tags added in the first PCR on the same molecule to detect strand exchange; as described in the data analysis, the rate of strand exchange was low presumably because we carefully limited the number of PCR cycles.

#### Deep mutational scanning for mutational effects on cell entry

To measure effects of G mutations on cell entry in 293T cells, we infected 293T cells with the rabies G pseudovirus library. To correct for library composition bias, we simultaneously infected a separate set of 293T cells with VSV-G expressing library pseudovirus (Fig S2A). Because VSV-G drives infection in the latter case, all pseudoviruses regardless of rabies G mutant function can mediate cell entry when VSV-G is present.

For all infection assays, we coated 6-well plates (Corning, Ref. No. 3516) with poly-L-lysine (Sigma-Aldrich, Ref. No. P4707-50mL). To do this, we added 1mL of poly-L-lysine to all plate wells and ensured full coating. After a 5 minute incubation, poly-L-lysine was aspirated, wells were washed with molecular biology grade water and allowed to dry at room temperature. We seeded each well with between 700,000 to 800,000 293T cells. Approximately 24 hours later after seeding cells, infection assays were performed.

For the rabies G pseudovirus library, we wanted each replicate well for the infection assays to have over 10-fold variant coverage in our library to prevent bottlenecking, so we infected with >1 million transducing units of virus per replicate well. (Note that the actual number of sequenced variants that enter cells may be higher, as transduction units are determined by flow cytometry which measured integrated viral genomes, but the non-integrated lentiviral DNA we mini-prep for sequencing is generally in substantial excess to integrated viral genomes.^92–94^) For our infection assays, rabies G pseudovirus library was first exchanged into D10 media adjusted to a pH of 7.1 and supplemented with 4µg/mL polybrene. To do this, we added 2 times the volume of pH 7.1 media to the virus used in a particular experiment. This was then spin concentrated at 3100*g* for 30 minutes using Amicon Ultra 15mL/100kD MWCO Centrifugal filters (EMD Millipore, Ref. No. UFC9100024). Total virus was then resuspended to a volume in pH 7.1 D10 media with polybrene so that each well would receive 500µL virus. To pH equilibrate media prior to infection, we combined virus with 600µL of pH 7.1 tissue culture media and incubated at 37°C and 5% CO_2_ in a tissue culture incubator. We also aspirated all media from cells and added 1mL of the pH 7.1 tissue culture media. Both virus and cells were incubated separately for 60 minutes, after which 1mL of virus was added to the cells. Infection assays proceeded for ∼12-14 hours.

For VSV-G expressing pseudovirus, we aimed for 100-fold coverage of barcoded variants and generally aimed to infect cells with ∼12-15 million transducing units of virus. Due to this high number, we split VSV-G pseudovirus infections across two wells of 293T cells such that each well was infected with ∼6 to 7.5 million transducing units. To ensure that infections occurred at an optimal pH for VSV-G fusion activity, we conducted these infections in tetracyline-negative standard D10 media (pH 7.4). VSV-G pseudovirus was also spin concentrated and resuspended in D10 media to a sufficient volume for each well of cells to receive 1 mL of virus. As for rabies G library pseudovirus, infections proceeded for 12-14 hours.

After the infection assays, cells were miniprepped to collect non-integrated reverse-transcribed lentiviral genomes using a QIAprep Spin Miniprep Kit (Qiagen, Ref. No. 2710). To prepare cells for miniprepping, media was aspirated from infected cells, and cells were trypsinized (Fisher, Ref. No. MT25053CI) and collected into microcentrifuge tubes. We washed cells with 1X PBS and miniprepped the cell pellet. To increase DNA yield, we warmed miniprep columns with 35µL of elution buffer added to 55°C for 5 minutes prior to spin-down elution.

#### Deep mutational scanning for mutational effects on antibody escape

The experiments to measure the effects of G mutations on escape from antibody neutralization were performed largely as above with minor alterations. Firstly, to convert sequencing counts into absolute neutralization, we used a “neutralization standard” consisting of a barcoded pseudovirus expressing VSV-G as described previously^36,37^. These pseudoviruses contain a defined set of barcodes and act as a non-neutralized standard for assays. This standard virus was added to the rabies G pseudovirus library at 1-2% of the rabies G pseudovirus library titer. As above, we used spin concentration to remove D10 media and to exchange with D10 media which has a pH of 7.1.

We performed infection assays at multiple concentrations of antibody to achieve levels of neutralization ranging from ∼50% to >99% as determined by the sequencing and comparison to the neutralization standard as described below. We combined 500µL of resuspended virus with 600µL of media supplemented with antibody so that the combined 1.1mL mixture contained antibody at the appropriate concentrations. We allowed antibody binding and pH equilibration to occur for 1 hour at 37°C and 5% CO_2_ in a tissue culture incubator as the cells in 1mL of pH 7.1 media. After this incubation period, virus was added to cells and infections proceeded for 12-14 hours. Infected cells were washed and non-integrated lentiviral DNA was miniprepped as above for cell entry experiments.

#### Illumina sequencing of pseudovirus barcodes

We used Illumina sequencing of the pseudovirus barcodes to quantify infection by each G variant in each condition. To prepare samples for sequencing, we employed a protocol involving two steps of PCR as described in Dadonaite et al.^36^ We used the first round of PCR to amplify barcodes and to append Illumina Trueseq Read 1 and Read 2 sequences. An Illumina Truseq Read 1 sequence was added via the forward primer annealing upstream of the barcode (Illumina_Rnd1_For), and the Read 2 sequence was added using the reverse primer annealing downstream of the barcode (Illumina_Rnd1_Rev). See https://github.com/dms-vep/RABV_Pasteur_G_DMS/blob/main/Additional_Data/PrimerSequences/RABV-G_Sequencing-Primers.fasta for sequences of the Illumina primers.

For each PCR, we combined 25 or 28µL of miniprepped non-integrated lentiviral DNA, 1.5µL of each primer (at 10µM stock), and 25µL of KOD Hot Start master mix. PCR cycling conditions were as follows:

1. 95°C, 2 min
2. 95°C, 20 sec
3. 70°C, 1 sec
4. 58°C, 10 s, cooling at 0.5°C/sec
5. 70°C, 20 sec
6. Return to Step 2, 27 cycles
7. 70°C, 1 min
8. 4°C hold

We cleaned PCR products with 150µL of Ampure XP beads each (3:1 volume ratio of beads to PCR reaction) and eluted in 50µL of elution buffer. We quantified concentrations of the PCR product with a Qubit 4 Fluorometer (ThermoFisher, Ref. No. Q33238).

For the second step of the PCR, we appended the P5 Illumina adapter overhang using a forward primer that anneals to the Illumina Truseq Read 1 sequence (IIllumina_Rnd2_For). The reverse primer annealed to the Truseq Read 2 sequence and contained the P7 Illumina adapter and i7 sample index (IIllumina_Rnd2_Rev). Indices were from the NextFlex 8nt barcode set. See https://github.com/dms-vep/RABV_Pasteur_G_DMS/blob/main/Additional_Data/PrimerSequences/RABV-G_Sequencing-Primers.fasta for sequences of the round 2 primers. Note that since indexes can vary, we have only provided a generic template where the index is denoted with ‘n’ characters. For each reaction, we combined 20 ng of round 1 PCR product, 2µL of each primer (at 10µM each) and added molecular biology grade water to 25µL total. To this, we added 25µL of KOD Hot Start Mix. We used PCR cycling conditions the same as above, except only 20 cycles total were performed.

We quantified round 2 PCR product concentration using a Qubit 4 Fluorometer (ThermoFisher, Ref. No. Q33238). We then pooled samples at masses (ng) proportional to the reads desired and ran the pool on a 1% agarose gel and extracted the band of the correct size (283bp). We ampure-cleaned the gel purified fragment and diluted the sequencing pool in elution buffer to a concentration of 4nM. Prior to sequencing, we confirmed the final sample contained a monodisperse peak around the expected product size via TapeStation.

We aimed to have at least 20 million single-end Illumina reads per barcoded sample in a pooled submission. For VSV-G library pseudovirus, we aimed for over 50 million reads per sample to ensure adequate depth for calculating library composition.

#### Production of luciferase-encoding pseudovirus for validation assays

We generated pseudovirus expressing rabies G and encoding luciferase for validation assays of how specific G mutations affected cell entry or antibody neutralization. Briefly, we seeded 750,000 293T cells into each well of 6-well plates. For each well, we transfected 250ng of G expression plasmid, 250ng each of 26_HDM-Hgpm2 (Gagpol), 27_HDM-tat1b (Tat), and 28_pRC-CMV-Rev1b (Rev) expression plasmids, and 1µg of a plasmid containing a lentiviral genome encoding luciferase and driven by a chimeric CMV-LTR promoter. Virus was harvested and sterile filtered via syringe (Corning, Ref. No. 431220). We generated all viruses in standard D10 at a pH of 7.4.

To titrate these pseudoviruses, we seeded 15,000 293T cells per well in black-wall, clear-bottom, poly-l-Lysine coated 96-well plates 24 hours prior to infections. On the day of infection, media was removed and replaced with 50µL of fresh D10 at pH 7.1 supplemented with 4µg/mL polybrene. We generated serial dilution courses of virus in separate plates proceeding in 2- or 4-fold dilution steps. This allows us to sample RLU values at multiple volumes to gather technical replicates. Because the pH 7.1 D10 media requires pre-equilibration in tissue culture incubator settings, we pre-equilibrated virus and cells separately for 60 minutes. We then added 100µL to the cells for infections.

After 48 hours, we measured RLU using the Bright-Glo Luciferase Assay System (Promega, Ref. No. E2620). Briefly, 100µL of media was aspirated from wells, and 30µL of Bright-Glo was added for a 1:1 volume ratio in each well. Cells were lysed and luciferase activity was measured by plate-reader. To reduce background luminescence, we placed a black sticker surface to mask the clear bottoms of all wells. We normalized counts to volume to generate RLU/µL to calculate volume required for 250,000-500,000 RLU for neutralization curves. For each virus, we calculated an RLU/µL from technical replicates by taking the mean of all individual replicates.

#### Validation pseudovirus neutralization assays

For neutralization assays to validate the effects of specific G mutations, we seeded 15,000 293T cells per well in black-wall, clear-bottom, poly-l-Lysine coated 96-well plates 24 hours prior to infections. On the day of infection, media was removed and replaced with 50µL of fresh D10 at pH 7.1 supplemented with 4µg/mL polybrene. We generated an antibody dilution course plate using 5-fold dilutions steps down from a maximum concentration. Each plate had 2 column replicates for each antibody, as well as 2 column replicates for wells having no antibody to measure initial RLU per well. We also had a reserved column with no virus as an autofluorescence control.

We diluted the virus into D10 at pH 7.1 supplemented with 4µg/mL polybrene so that a volume of 60µL would yield approximately 250,000-500,000 RLU. To a new plate, we added 60µL to each well (except for those serving as a no-virus control). We combined 60µL from each well in the antibody plate and allowed binding and pH equilibration to proceed for 60 minutes. Following this, we took 100µL from each well and added them to the plate containing cells.

RLU for neutralization assays were measured as for luciferase titer assays. RLU values in each plate were mapped to a virus, antibody identity, and antibody concentration. We also took the median of luminescence values and subtracted each RLU measurement to correct for plate background. To calculate fraction infectivity, we calculated the median RLU of all wells containing no antibody and normalized all RLU measurements to this quantity. We fit neutralization curves to the fraction infectivities using the neutcurve package^95^ (https://jbloomlab.github.io/neutcurve/index.html).

#### Pseudovirus validations of mutational effects on cell entry and antibody escape

To validate cell entry and antibody escape mutations in rabies G, we generated individually cloned luciferase viruses that express glycoprotein mutants. To do this, we first cloned point mutations in an expression plasmid encoding the Pasteur strain of rabies G (https://github.com/dms-vep/RABV_Pasteur_G_DMS/blob/main/Additional_Data/PlasmidSequences/3968_HDM_Rabies-G_Pasteur_GenscriptOpt3.gb). Mutagenesis primers were generated using the NEBaseChanger tool. See (https://github.com/dms-vep/RABV_Pasteur_G_DMS/blob/main/Additional_Data/PrimerSequences/RABV-G_Mutagenesis-Primers.fasta) for all of the mutagenesis primers. For each mutagenesis reaction, we combined 10ng of unmutated expression plasmid, 1.25µL each of corresponding forward and reverse primers (10µM stock), and 12.5µL of Q5 High-Fidelity 2X Master Mix (New England Biolabs, Ref. No. M0492S). We used the kit’s standard PCR cycling conditions and suggested annealing temperatures. To digest unmutated template plasmid and ligate PCR products, the PCR products were treated with KLD Enzyme Mix (New England Biolabs, Ref. No. M0554S). We transformed a small volume of the PCR product into Stellar Chemically Competent cells (Takara, Ref. No. 636766). These expression plasmids were used as before to make luciferase-encoding pseudovirus (see “Production of luciferase-encoding pseudovirus” section). See https://github.com/dms-vep/RABV_Pasteur_G_DMS/blob/main/Additional_Data/PlasmidSequences/4954_HDM_Rabies-G_Pasteur_I338T.gb for an example mutant plasmid map.

We performed titrations using this virus as described above (see “Titration of virus via luciferase readout”). To compare with functional effects from the deep mutational scan, we normalized these titers to those of unmutated Pasteur rabies G pseudovirus (Fig 1D). For glycoprotein mutants with entry scores below −5 (H270P and N319A), we set cell entry to −5 as this was equivalent to the effect of stop codons. We performed replicates using independent stocks of miniprepped expression plasmids. See https://github.com/dms-vep/RABV_Pasteur_G_DMS/blob/main/non-pipeline_analyses/Additional_Notebooks/240810_LuciferaseValidations.ipynb for raw data and notebooks for analysis.

The validation assays for escape mutations from antibody 17C7 shown in Fig S3D were performed and analyzed as above in the “Neutralization assays” section with minor changes. Firstly, we aimed for a maximum antibody concentration of 5µg/mL. Secondly, we adjusted the layout of the plates to accommodate the necessity of no antibody controls for 4 separate viruses per plate. See https://github.com/dms-vep/RABV_Pasteur_G_DMS/blob/main/non-pipeline_analyses/Additional_Notebooks/17C7_EscapeValidations.ipynb for analysis.

#### Validation of deep mutational scanning predictions using natural rabies G strains

We identified rabies G protein sequences for two circulating strains A12_2718 (NCBI Accession No. KC792123.1) and NY-2011-1202O (NCBI Accession No. QEJ74619.1) and codon sequence optimized using software from Twist or Genscript. We ordered these fragments from Twist and cloned them into the expression vector using HiFi Assembly. See https://github.com/dms-vep/RABV_Pasteur_G_DMS/tree/main/Additional_Data/PlasmidSequences for plasmid maps for the wild-type glycoproteins and for mutants generated by QuickChange (described below). The plasmid encoding the wild-type A12_2718 glycoprotein is 4956_HDM_Rabies-G_AGN94399-WT, and the plasmid encoding the wild-type NY-2011-1202O glycoprotein is 4959_HDM_Rabies-G_QEJ74619_WT.

Strains were pseudotyped onto lentivirus and titered for luciferase activity as above with minor modifications. For producing pseudovirus expressing the NY-2011-1202O glycoprotein, we produced virus at a scale of ∼60mL and concentrated via centrifugation into 1mL of virus to generate sufficient titers. We also performed plate-reader measurements in white-walled plates to reduce the amount of virus required.

To study the contribution of individual mutations in the A12_2718 background, we made mutations using the QuickChange protocol as described in the “Pseudovirus validations of mutational effects on cell entry and antibody escape” section. Virus was rescued and concentrated for the T338I mutant glycoprotein. To ensure sufficient titers for neutralization assays, we also measured titers in white-wall plates. Neutralization assays for antibodies were performed as previously described in the “Neutralization assays” section. With the exception of the experiment measuring neutralization of unmutated A12_2718 rabies G by 17C7 and RVC58 (shown in Fig 6C), we transferred the luciferase and cell mixture into white-wall plates for measurements for sufficient luciferase signal. Neutralization curves were subsequently processed using the neutcurve package as above (see “Neutralization Assays”).

### Computational Methods

#### PacBio sequencing analysis

To analyze PacBio sequencing data to link the barcodes to the G mutant in the lentiviral backbones, we used the *alignparse* package (https://github.com/jbloomlab/alignparse)^96^. We subsequently filtered sequencing reads on various criteria. Firstly, we removed Circular Consensus Sequencing reads (CCSs) that contained an error rate higher than 10^−4^. This removed ∼25% of reads for Libraries A and B. We also used the G or C nucleotide tags added during PCR to remove strand-exchange sequences, which removed ∼1% of sequences per library (validating that the rate of strand exchange during the library-prep PCR was low). These CCSs were then aligned to a rabies G reference sequence, and we generated consensus sequences for each barcode requiring at least 3 CCSs and below 20% of minor sub or indel frequencies. We generated a barcode to variant look-up table with these sequences with an empirical accuracy of ∼0.8 for both libraries. After this process, Library A contained 83,706 variants, and Library B contained 89,327 variants (Fig S1C).

See (https://github.com/dms-vep/RABV_Pasteur_G_DMS/blob/main/results/variants/codon_variants.csv) for the final barcode to variant table. Jupyter notebooks are available with the code for analyzing PacBio CCSs (https://dms-vep.org/RABV_Pasteur_G_DMS/notebooks/analyze_pacbio_ccs.html), building consensus sequences (https://dms-vep.org/RABV_Pasteur_G_DMS/notebooks/build_pacbio_consensus.html) and building the final barcode-variant table (https://dms-vep.org/RABV_Pasteur_G_DMS/notebooks/build_codon_variants.html).

#### Illumina barcode sequencing analysis

We used the tool illuminabarcodeparser (https://jbloomlab.github.io/dms_variants/dms_variants.illuminabarcodeparser.html) to count barcodes mapping to each variant for each experimental conditions. We filtered for valid barcodes by examining nucleotide sequences downstream of a standard sequence in the amplicon with only 2 mismatches allowed. Barcodes were required to have a minimum quality score of 20. For each experiment, we required that barcodes met a minimum count threshold in “pre-selection” conditions, either VSV-G pseudovirus barcode counts in functional effect selections or no-antibody conditions in antibody escape assays. This step removes barcodes that are poorly represented in specific experiments. The specific thresholds used for each experiment is specified in the YAML configuration files on the GitHub repo (https://github.com/dms-vep/RABV_Pasteur_G_DMS).

#### Calculation of effects of mutations on cell entry

We first assigned a cell entry functional score to each barcoded variant^36,37^. Briefly, for a barcode mutant *m*, we calculated functional scores using the formula 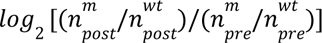, where 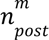 is the barcode count for a mutant in the rabies G expressing pseudovirus library, 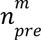 is the barcode count for a mutant in the VSV-G expressing library, 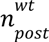 and 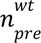 are the corresponding counts for all unmutated (wildtype) G barcoded variants in the library. Therefore, G mutants that have wildtype-like cell entry will have a functional score of zero, while mutants with impaired G entry will have functional scores less than zero. Because some mutants in the library contain multiple amino acid mutations, we applied a global epistasis model^40^ with a sigmoid global epistasis function as implemented in *multidms*^97^ to deconvolve the effects of individual amino acid mutations. In this paper and in the interactive displays at https://dms-vep.org/RABV_Pasteur_G_DMS/cell_entry.html we report the average cell entry effects for mutations across libraries and experiments. Basic filtering for high-confidence measurements was applied by only reporting values for mutations seen in an average of at least two barcoded variants per library and with measurements in the majority of experimental replicates.

#### Calculation of effects of mutations on antibody escape

To calculate the effects of mutations on antibody escape, for each barcoded variant, we first determined the non-neutralized fraction at each antibody concentration by comparing the barcode counts for the variant to those for the non-neutralized VSV-G standard barcodes as previously described in Dadonaite et al.^36^ We then used the *polyclonal*^45^ software package (https://jbloomlab.github.io/polyclonal/) to estimate the effects of each mutation on escape. The mutation effects on escape are linearly correlated with the change in log IC50 caused by each mutation, with positive values indicating escape and negative values indicating increased neutralization. See the “Antibody/serum escape” section of https://dms-vep.org/RABV_Pasteur_G_DMS/appendix.html for notebooks with details about the fitting and that show the fraction of the library that is non-neutralized at each antibody concentration.

For all figures and interactive visualizations (https://dms-vep.org/RABV_Pasteur_G_DMS/escape.html), we applied filters to remove low-confidence measurements. Specifically, we only showed mutations observed in an average of at least two-barcoded variants across libraries, and by default we hide the escape effects for mutations that are highly deleterious for cell entry (as it is difficult to measure the neutralization of G variants that are largely non-functional). The default visualizations also floor escape values at zero (so only show positive escape). All of these options can be adjusted using the interactive visualizations at https://dms-vep.org/RABV_Pasteur_G_DMS/escape.html.

#### Nextstrain phylogenetic trees of publicly available rabies G sequences

We gathered publicly available rabies G sequences from NCBI Virus using the query term “Lyssavirus rabies, taxid:11292”in the “Search by virus name or taxonomy box.” Sequences were downloaded as of October 18, 2024. Only G sequences with at least 1500 nucleotides and no ambiguous nucleotides (“N”) were included. The G sequences were processed and aligned using Augur^98^, and a phylogenetic tree was inferred from the resulting alignment. The phylogenetic tree was rooted using the non-rabies lyssavirus (also in phylogroup I) Gannoruwa bat lyssavirus (NCBI accession: NC_031988) as an outgroup.

Antibody escape scores were mapped onto the tree by summing the escape measurements of G sequence substitutions relative to the Pasteur G sequence, and the tree was visualized using Nextstrain^68^. See https://nextstrain.org/groups/jbloomlab/dms/rabies-G for the interactive Nextstrain trees, and https://github.com/dms-vep/RABV_Pasteur_G_DMS/tree/main/non-pipeline_analyses/RABV_nextstrain for the computer code used to generate the trees. Note that the trees can be colored by G amino-acid identity using either the G ectodomain numbering scheme used in this paper, or sequential numbering of the entire G protein.

Note that the Nextstrain phylogenetic trees use all 7,122 publicly available sequences, but the plot in Fig 6A uses only the 6,581 sequences corresponding to natural sequences (excluding lab-passaged ones and those originating from an unknown host).

#### Structural analysis

We examined the effects of mutations on several previously determined structures of rabies G. These structures include pre-fusion conformation (PDB: 7U9G), extended intermediate conformation (PDB: 6LGW) and rabies G in complex with antibodies (PDB 8R40 for CR57, PDB 8A1E for 17C7, PDB 7U9G for RVA122, and PDB 6TOU for RVC20). To project the mutation effects on structures, we made a notebook (https://github.com/dms-vep/RABV_Pasteur_G_DMS/blob/main/non-pipeline_analyses/Additional_Notebooks/color_pdb_functionaleffects.ipynb) that takes a PDB file, relevant chain IDs, and a CSV of deep mutational scan measurements and re-assigns the b-factors in the PDB file to the desired measurements. For cell-entry we used a site statistic of the mean effect of mutations at each site; for antibody escape we used the summed positive escape values across all mutations at that site.

See https://github.com/dms-vep/RABV_Pasteur_G_DMS/blob/main/non-pipeline_analyses/Additional_Notebooks/241126_GetContacts.ipynb for jupyter notebooks used to determine all residues within 4Å of an antibody in complex structures and producing heatmaps.

